# Lateral entorhinal cortex subpopulations represent experiential epochs surrounding reward

**DOI:** 10.1101/2023.10.09.561557

**Authors:** John B. Issa, Brad A. Radvansky, Feng Xuan, Daniel A. Dombeck

**Affiliations:** Department of Neurobiology, Northwestern University, Evanston, IL 60608, USA

## Abstract

During goal-directed navigation, “what” information, which describes the experiences occurring in periods surrounding a reward, can be combined with spatial “where” information to guide behavior and form episodic memories^1,2^. This integrative process is thought to occur in the hippocampus^3^, which receives spatial information from the medial entorhinal cortex (MEC)^4^; however, the source of the “what” information and how it is represented is largely unknown. Here, by establishing a novel imaging method, we show that the lateral entorhinal cortex (LEC) of mice represents key experiential epochs during a reward-based navigation task. We discover a population of neurons that signals goal approach and a separate population of neurons that signals goal departure. A third population of neurons signals reward consumption. When reward location is moved, these populations immediately shift their respective representations of each experiential epoch relative to reward, while optogenetic inhibition of LEC disrupts learning of the new reward location. Together, these results indicate the LEC provides a stable code of experiential epochs surrounding and including reward consumption, providing reward-centric information to contextualize the spatial information carried by the MEC. Such parallel representations are well-suited for generating episodic memories of rewarding experiences and guiding flexible and efficient goal-directed navigation^5–7^.

## Introduction

Episodic memories of rewarding experiences include information about where episodes occurred and also non-spatial information about what occurred (event context)^8^. For example, a dinner out is an experience that consists of clear spatial navigation components but also different experiential epochs: experiencing hunger and anticipation before arriving at the restaurant, consuming dinner and experiencing the taste of the food, experiencing contentment and satiation after departing the restaurant. These experiential epochs are not obviously aligned to sensory inputs but are important components in forming episodic memories. Here we refer to such non-spatial components of these epochs as reward experience epochs.

The integration of spatial and experiential information is thought to occur in the hippocampus^3^, where place cells encode not just locations in an environment but also additional non-spatial information^9,10^, as evidenced by the context-dependent firing of place cells during choice tasks^11,12^, in response to aversive stimuli^13^, and more^14,15^. This context-dependence of place cells is especially prominent in reward-guided navigation, which leads to an overrepresentation in the number of place cells encoding the region around reward^16,17^ and distinct signaling of goal approach and goal departure^18,19^. How the “what” information about reward experience reaches the hippocampus is poorly understood, however. In particular, such information should not just encode the discovery of a reward but also represent information from the periods leading up to reward and after reward (such as of the dinner out; Figure 1a)^1,2,20^.

**Figure 1:**
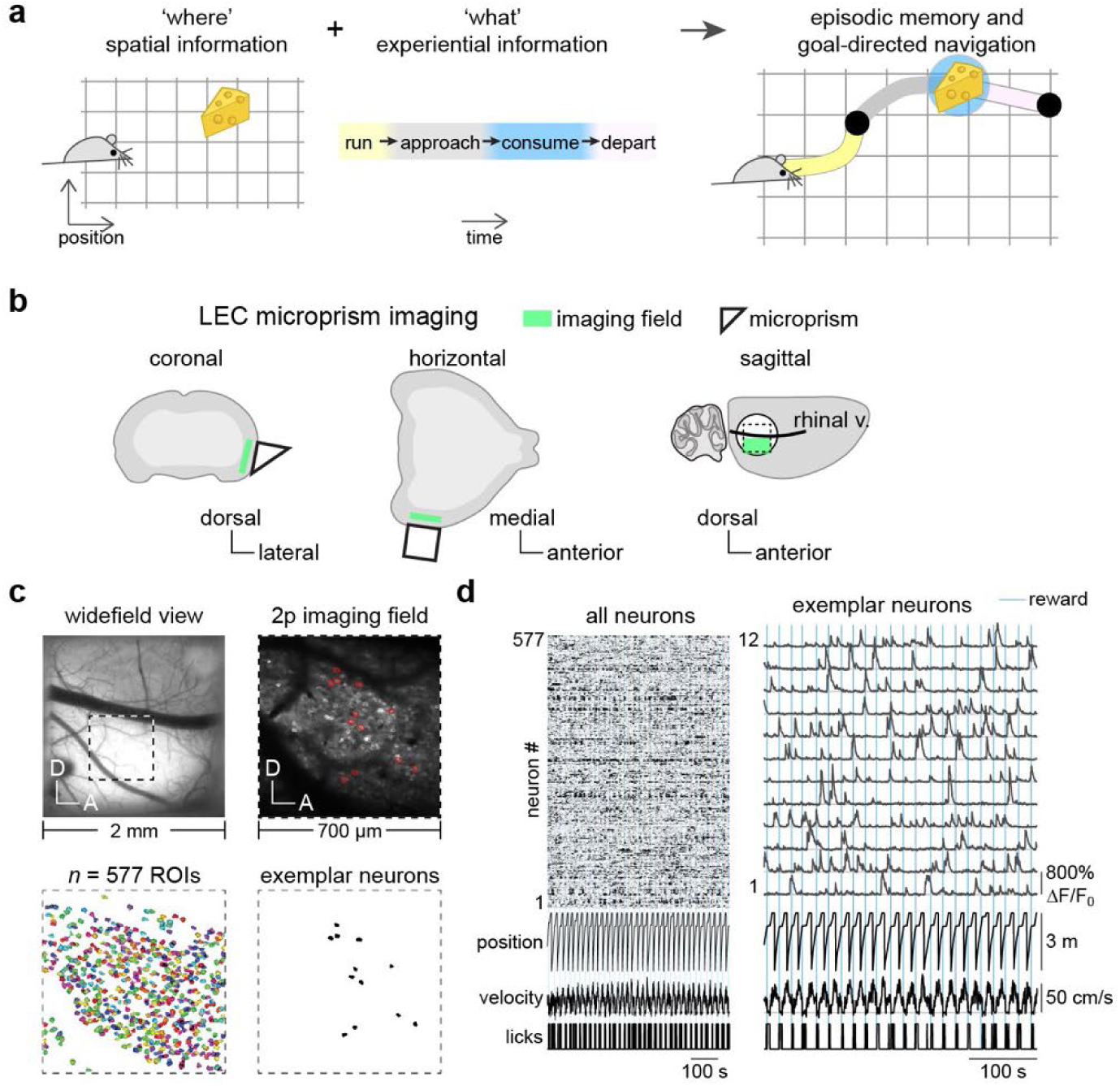
Two-photon imaging of the lateral entorhinal cortex. a) A schematic of goal-directed navigation that requires a spatial map along with a reward experience representation. In this proposed model, episodic memories of goal-directed navigation combine spatial “where” information with experiential “what” information during each epoch of the reward experience. b) Schematic of LEC imaging preparation from three views. Green region denotes targeted region. Circle represents 3 mm No. 0 glass coverslip; square/triangle (2×2 mm) represents 45-degree microprism. c) Views of exemplar field as seen through microprism using widefield and two-photon fluorescence imaging of neurons expressing GCaMP6s. Underneath is shown segmentation of individual cells and cell masks for all 577 neurons and 12 exemplar neurons to be shown in panel d. d) Fluorescence traces for all detected neurons (clustered using k-means to aid visualization) and expanded view for 12 exemplar neurons from panel c after neuropil subtraction and baseline correction. Behavioral signals are shown underneath.

The medial entorhinal cortex (MEC), a major source of cortical input to the hippocampus^21^, contains spatially modulated cells such as grid cells^22^, border cells^23^, and head direction cells^24^, which convey spatial information to the hippocampus. In addition to these spatial representations, the MEC could also be a potential source of reward experience information to the hippocampus. As such, multiple studies have investigated the role of the MEC in reward-guided navigation^25–27^. Bulk MEC activity is only moderately influence by the presence of rewards^27^ and, while a rewarded location could distort nearby firing fields of grid cells, these changes were not shown to be specific to the trajectory^25^. Thus, MEC has not been shown to represent different reward experience epochs^27^.

Alternatively, neuromodulatory systems could provide such reward experience information to the hippocampus^28^. Most notably, the locus coeruleus (LC) provides bursts of noradrenaline and dopamine to the hippocampus at a new reward location during navigation^29,30^. Inhibition of these inputs prevented the formation of overrepresentation of reward locations by place cells. Importantly, however, activation of these inputs alone was not sufficient to drive the overrepresentation, as concurrent rewards were still required^29^. This result indicates that LC inputs to the hippocampus provide a learning signal that can open a plasticity window for spatial information to be associated with reward experience information but, crucially, the reward experience information appears to reach the hippocampus from a separate and currently unknown source.

Finally, the lateral entorhinal cortex (LEC), which similar to the MEC provides significant excitatory input to the hippocampus^21^, could possibly provide the reward experience information. While previous studies have uncovered a variety of roles for the LEC across a range of behaviors, including object coding^31,32^, timing^33^, olfaction^34^, and nonspatial associative learning^35^, its role in representing rewards and reward experience has not been directly addressed. Critically, prior studies did not disentangle rewards from objects, making it unclear whether LEC was encoding the object salience itself or valence related to the reward experience. Furthermore, large-scale single neuron resolution recording has proven challenging in this difficult-to-access brain region, hindering investigation of the myriad functions supported by LEC. To overcome these challenges, here we developed a novel two-photon imaging technique that allowed for simultaneous functional recordings of an unprecedented number of LEC neurons. We also made use of a virtual reward-guided navigation task, which added valence to an otherwise unmarked location, thus dissociating object salience and reward experience.

## Results

### Two-photon imaging of the lateral entorhinal cortex in behaving mice

Two-photon functional imaging offers the capability to record from large numbers of neurons simultaneously, but this technique has not previously been used in the LEC during behavior^34^. Therefore, we first sought to develop a method for two-photon imaging of the LEC in behaving mice. As it is situated ventral to the rhinal fissure^36^, the LEC is a lateralized structure in rodents, and direct access with a microscope requires approaching from an angle greater than 90 degrees to the horizontal plane. A further complication is the surrounding anatomy: the pinna, nearby vasculature such as the petrosal squamosal sinus, and protrusions of the skull (zygomatic process) each impinge upon optical access with a microscope^37^. To overcome these limitations, we developed surgical methods to implant a cranial window (3 mm round coverslip) with an attached microprism (2.0 mm square) to rotate the imaging plane 90 degrees (Figure 1b). Using this approach in transgenic mice expressing the fluorescent Ca^2+^ indicator GCaMP6s^38,39^, we could capture a large field of view of the LEC using a conventional upright two-photon microscope and access depths > 250 µm (Figure 1c) in head-fixed mice running on a treadmill to traverse a 1-D virtual track. Movies were motion corrected and cells identified and segmented using Suite2p^40^. Neuropil-corrected, baseline-adjusted, and deconvolved neural Ca^2+^ transients (referred to here as “firing”) were recovered using a novel iterative algorithm (Methods). In a typical imaging field of 700 x 700 µm, ~500 active cells were observed during behavior (mean across 47 imaging fields: 496 cells; range: 150 to 843 cells) (Figure 1d). Having established a new method to record from large populations of individual LEC neurons in behaving mice, we next studied the encoding properties of LEC populations during reward-guided navigation.

### Enrichment of LEC firing near reward locations segregates into pre- and post-reward populations

In order to examine how the LEC may represent reward experience epochs, we designed our virtual track to be visually cue rich but the reward location itself not marked by any visual feature or object. This design helped isolate visual object coding from reward coding, which we exploit in later sections to move the reward without changing other aspects of the environment. Mice learned over several training sessions to decelerate and lick in anticipation of the reward location, where a drop of water was delivered (Figure 2a). We began by examining the firing of neurons in LEC along the track during this behavior since reward experience is expected to change along the track, leading to spatial information as an initial means to identify reward experience encoding neurons. In parallel, we also performed two-photon Ca^2+^ imaging of populations of neurons in MEC and in CA1 in separate groups of mice implanted with microprisms^41^ or cannulas^42^, respectively (Figure 2b). Analyses of changes in reward location are left to later sections; here, we examine coding of a learned reward location. Three different reward locations were used in our experiments. We first show data from the most commonly used reward location (2.3 m) and pool data from the other reward locations (0.7 m and 1.5 m) where appropriate.

**Figure 2:**
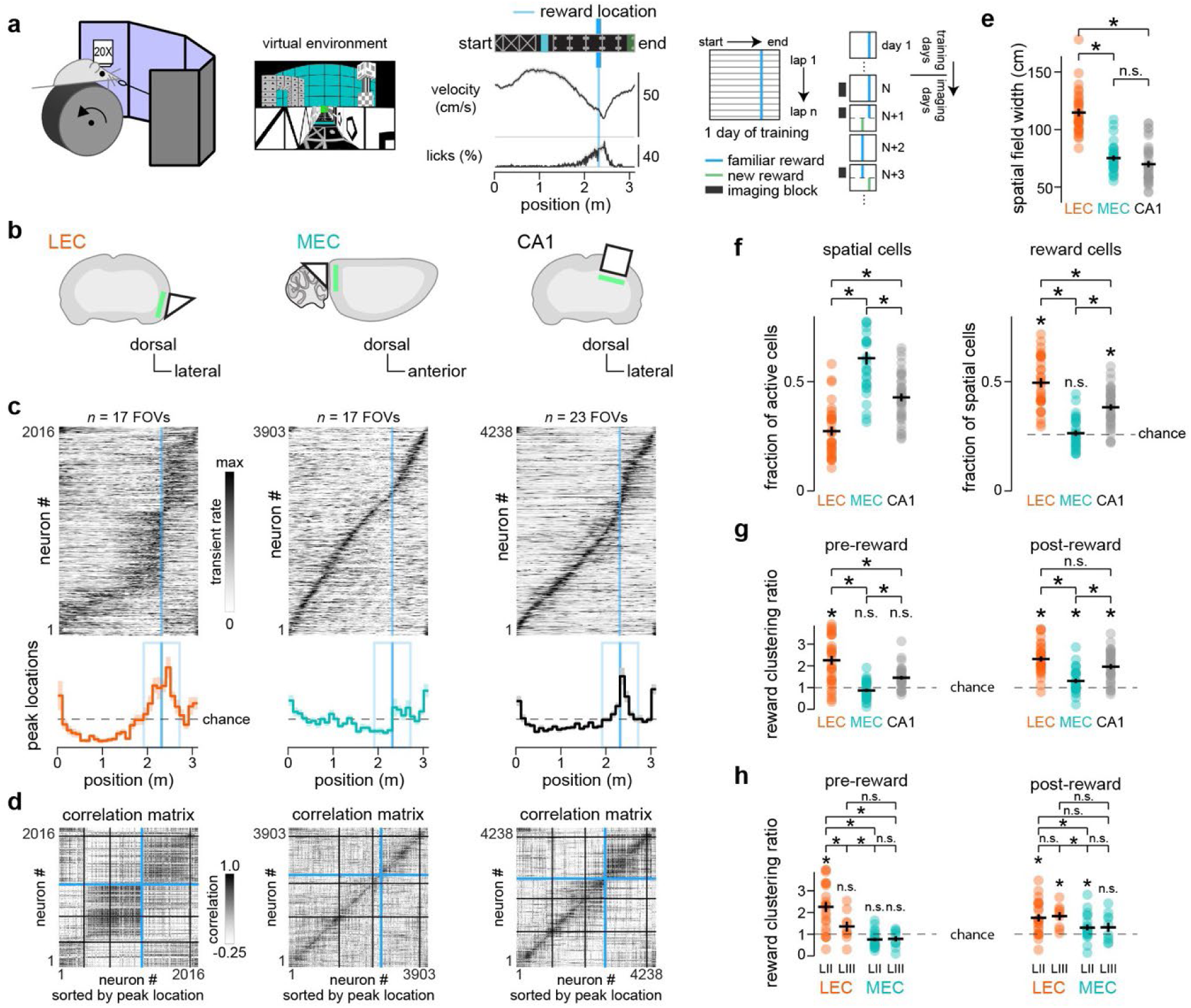
Prominent reward clustering in LEC with segregation of pre- and post-reward populations. a) Head-fixed mice traverse a 3.1 m linear track in virtual reality with water reward delivered at 2.3 m. Treadmill velocity and detected licks, averaged over 43 traversals, are binned at 1 cm intervals with stationary periods excluded (Methods). Training paradigm shown to the right. Once behavior reached criteria (at least two laps per minute over a ~40 minute training session along with the presence of anticipatory licking and deceleration for reward), imaging sessions began. On subsequent days, reward location was moved to either 0.7 m or 1.5 m. At least one session with the new reward location was interspersed to allow for it to become familiar to the animal as assessed by anticipatory behavior of the new reward location. b) Schematics of imaging approaches using implanted microprisms (LEC, MEC) or cannulae (CA1). c) Population spatial firing patterns for LEC, MEC, and CA1 mice. Spatial cells, defined as neurons with significant spatial information (Methods), are sorted by where their firing peaked along the track on even laps. Each row of the image is the firing of one neuron averaged over odd laps, binned at 1 cm intervals after Gaussian filtering (170 ms standard deviation), and normalized to its maximum value. Histograms underneath each plot show proportion of peak locations for neurons in 10-cm bins, with ‘chance’ calculated as expected number in each bin if uniformly distributed (1/31 bins) and ‘reward cells’ those that peak within the blue rectangle (±40 cm from reward). Histogram is calculated for individual sessions (FOV, or field-of-view); mean ± SEM are shown as dark line with light shading, respectively. d) Correlation matrix formed from the sorted heat maps in panel B. Each pixel is the Pearson correlation coefficient between pairs of neurons. Sorting is same as in B. Blue lines indicate division between pre-reward and post-reward neurons. Black lines indicate 1, 2, and 3 m locations. Correlations are calculated between average firing on even and odd laps, so the diagonal is not necessarily equal to one. e) Quantification of spatial field widths in each region, computed as the distance of the track for which the firing rate is at least 30% of the maximum for each spatial cell. Each point represents the mean of spatial mean widths for all spatial cells from one imaging session; black cross represents mean ± SEM across sessions. Statistical tests performed between each pair of regions (2-sample t-test). f) Fraction of active cells that are spatial cells and fraction of spatial cells that are reward cells in each region. Each point represents one imaging session; black cross represents mean ± SEM across sessions. Statistical tests performed between each pair of regions (2-sample t-test) and, for reward cells, compared to chance as well (1-sample t-test). Chance was calculated assuming uniform distribution of spatial cell peaks along the track. Since the reward zone is 80 cm total and the track length in 310 cm, chance is 80/310, or ~0.26. g) Reward clustering ratio, calculated as the number of cells that peak in each reward zone (blue rectangle shown in c) divided by the expected number of cells if peaks were uniformly distributed along the track. Ratio greater than one (chance) indicates an increased number of cells in that reward zone. Statistical tests performed between each pair of regions (2-sample t-test) and compared to chance (1-sample t-test). h) Reward clustering ratio stratified by imaging depth. For LEC: up to 150 µm below dura (layer II) and at least 200 µm below dura (layer III); for MEC: up to 150 µm below dura (nominal layer II) and at least 150 µm below dura (nominal layer III). Statistical tests performed between each pair of regions (2-sample t-test) and compared to chance (1-sample t-test). For all statistical tests: * indicates p < 0.05 and n.s. indicates p > 0.05.

Among active neurons in the LEC, we selected for those with significant spatial information along the track^43^ (Methods). From 17 fields-of-view across 7 mice with the reward located at 2.3 m, 2016 such spatial cells were found, which represent 24.9% of active cells (Figure 2f). Across all reward positions, we identified 3956 spatial cells out of 14489 active neurons from 32 fields-of-view (Supplementary Tables 1 and 2). We plotted their spatial tuning, with neurons sorted by where their mean firing peaked along the track. Nearly half of spatial cells in LEC were active near the reward location, with such reward cells defined as spatial cells whose mean firing peaked within 40 cm of the reward location (Figure 2c,f). By comparison, MEC spatial cells were active nearly uniformly across the track, and the fraction active around reward was as expected by chance, while in CA1 an enhanced fraction of cells was active near reward, but less than what was observed in LEC (Figure 2c,f and Supplementary Figure 2a). Track position could still be decoded from activity in non-reward spatial cells in LEC, but the decoder performance error was higher for LEC than in MEC or CA1 (p < 0.05, 2-sample t-test; Supplementary Figure 2c). Cells from all three brain regions also tended to cluster near the start and end of the track, but we focused our analysis on the cells active around the reward location.

Beyond clustering near the reward, LEC spatial cells had wide spatial fields and were separated by whether they were active before or after the reward location (Figure 2c). These features are clearly visible when plotting the correlation matrix of the mean spatial firing maps, sorted by where each cell peaked (Figure 2d). While MEC shows a clear diagonal band structure, indicating narrow spatial fields across all track locations, LEC shows a block-like structure, indicating wide spatial fields (Figure 2e) that largely segregate into pre-reward-active neurons and post-reward-active neurons, a division confirmed by k-means clustering (Supplementary Figure 2d). To quantify the enrichment of spatial cells near reward, we divided the number of cells that peak in each reward zone (either pre- or post-reward) by the expected number of cells if peaks were uniformly distributed. This reward clustering ratio was enhanced in LEC for pre-reward compared to CA1 and MEC and in both LEC and CA1 for post-reward (Figure 2g).

The superficial layers of LEC are known to differ in cell type and connectivity^21^, so we next explored whether laminar differences may relate to reward clustering within LEC. In particular, layer II contains fan cells, which receive an outsized number of dopaminergic inputs from the VTA^35,44^, suggesting a potential region where reward information may enter the system. We imaged at depths targeting the two layers: 80-150 µm below dura for layer II (23 fields) and 200-250 µm below dura for layer III (9 fields) (Supplementary Figures 1 and 2b). While both regions show preferential firing near the reward location, the enhancement of firing in the pre-reward region was significantly larger in layer II versus layer III (Figure 2h). Importantly, such a difference did not exist for MEC, with a paucity of pre-reward neurons across imaging depths. To determine whether the differences in reward-related neural responses across LEC layers could be explained by other factors, we performed a multiway analysis of variance using imaging depth, age, gender, and deceleration (a behavioral measure of task performance) as predictors of reward clustering and found that only depth provided a significant prediction (Supplementary Table 1).

Therefore, we found that LEC, unlike MEC, was highly active around the reward location, with populations of neurons segregated by their firing during either goal approach or goal departure and the goal-approach population particularly enriched in layer II of LEC.

### Location-invariant representation of reward by pre- and post-reward cells in the LEC

We next asked whether the pre-reward and post-reward populations were dedicated to encoding particular epochs around reward irrespective of the reward’s location. To disentangle whether reward cells are encoding spatial information (track position) or reward itself, the reward location was moved without any cue, indication, or change to the environment in the middle of a session after ~40 laps with a familiar reward location. Mice learned to anticipate the new reward location by shifting their deceleration along the track (Figure 3a). Imaging was performed throughout the session, thus capturing representations both before and after the reward switch in the same cells.

**Figure 3:**
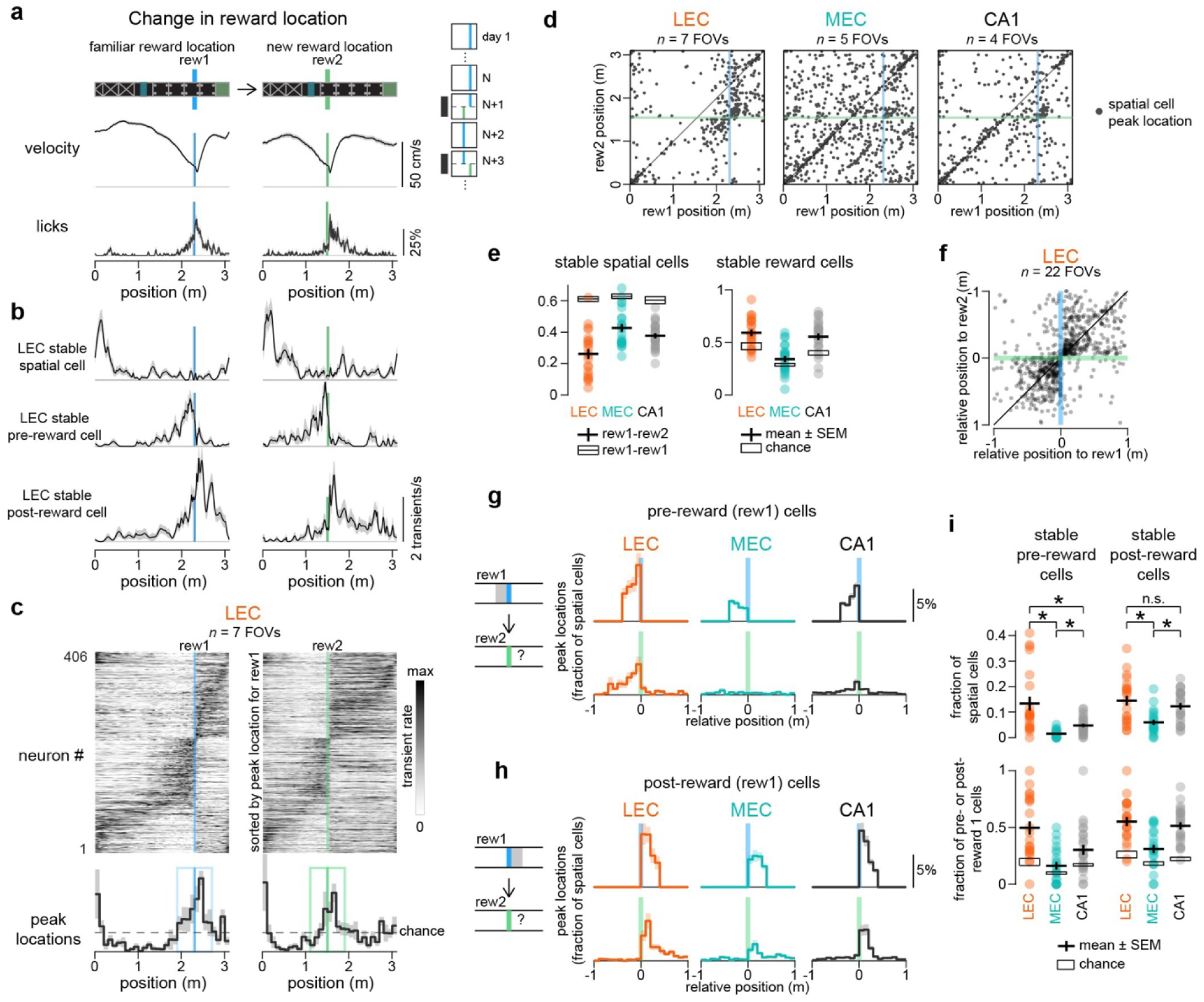
Dedication of pre- and post-reward populations in the LEC. a) Licking (percent of laps with contact of sensor) and velocity before (66 laps) and after (49 laps) change in reward location with ±SEM shown in light shading. In this example session, the reward was moved earlier on the track, from a familiar location (rew1, 2.3 m, blue) to a new location (rew2, 1.5 m, green). Inset shows training paradigm. Once a reward location became familiar, an imaging session was performed spanning ~15 minutes with the familiar reward location and ~15 minutes with the new reward location. b) Spatial firing (deconvolved Ca^2+^ transients) for three exemplar neurons from the same session as a, with ±SEM shown in light shading. Imaging was performed for a subset of laps (56 laps before and 44 laps after change in reward location). c) Spatial firing patterns along the track and histogram of firing peaks for spatial cells sorted by their firing peaks with the familiar reward location, analogous to Figure 2c. Same sorting is used to plot the firing of these cells with respect to the new reward location. Only cells that pass criteria for spatial information under both conditions are included. d) Dot raster of peaks for spatial cells in LEC, MEC, and CA1. Cells on the diagonal maintain their firing position along the track while cells at the intersection of reward locations (blue and green lines) adjusted their firing positions relative to the reward locations. e) Histograms measure the fraction of spatial cells that are stable after the reward location is moved, quantified by the peak location of the spatial tuning curve changing less than 40 cm (rew1-rew2). Data quantified for each imaging session; black cross represents mean ± SEM across sessions. For comparison, we computed the difference in peak location for even and odd trials with the original reward location (rew1-rew1); black rectangle represents mean ± SEM across sessions. Next, the fraction of reward cells that remain reward cells after the reward location change is quantified by the fraction of reward cells that maintain their peak location within 40 cm of the new reward location. f) Peak locations shown relative to reward locations with familiar reward condition on the x-axis and new reward condition on the y-axis. Each point represents one LEC neuron that was a spatial cell under both conditions. g) Peak locations for pre-reward cells relative to the familiar and novel reward locations. Selected cells are chosen for being pre-reward cells with the familiar reward location (rew1) and spatial cells in both; what is shown here is where their peak locations are for the novel reward location (rew2). Histograms are calculated for each imaging session; mean ± SEM are shown as dark line with light shading, respectively. h) Same as panel g but for post-reward cells. i) Histograms quantify cells that are stable pre- or post-reward cells at both reward locations, either as a fraction of all spatial cells or as a fraction of the pre- or post-reward cells for the familiar reward location. Each point represents one imaging session; black cross represents mean ± SEM across sessions; black rectangle represents mean ± SEM expected by chance. Statistical tests performed between each pair of regions (2-sample t-test). * indicates p < 0.05 and n.s. indicates p > 0.05.

Some LEC neurons maintained their firing field with respect to track position (LEC stable spatial cell, Figure 3b) and others with respect to the reward location (LEC stable pre- and post-reward cells, Figure 3b). We plotted all spatial neurons before and after this change in reward location from 2.3 m to 1.5 m, sorted by their firing peaks with respect to the familiar reward location (Figure 3c). Reward clustering again developed around the new reward location. Indeed, relatively few LEC neurons continued to fire at the same track location, especially compared to MEC and CA1 (Figure 3d-e, Supplementary Figure 3a-b). Instead, over half of the cells active in the reward zone for the familiar location shifted their firing fields to the new reward location (stable reward cells), more than expected by chance.

An even more significant picture of dedicated cell populations emerged when examining pre-reward and post-reward cells as distinct groups (Figure 3c,f). We found that nearly all pre-reward cells for the familiar reward location remained pre-reward cells relative to the new reward location and, similarly, post-reward cells remained post-reward relative to the new reward location (Figure 3f-h). The probability of a pre-reward cell remaining a pre-reward cell was 2.7x chance and 2.4x for post-reward cells (geometric mean of ratio for each session, Figure 3i). This dedication was largely absent in the MEC, as pre-reward cells for the familiar and new reward locations were not significantly related (Figure 3g,i) and the post-reward cells were only partially preserved (Figure 3h-i). Cells in CA1 showed a mixture of the strong dedication from LEC and the weak dedication from MEC (Figure 3g-i). When we teleported mice to new environments with unmarked reward locations, we found that reward dedication was similarly maintained in the new environment (1.5x for pre-reward and 2.3x for post-reward cells; Supplementary Figure 3c-h). Together, these results indicate that, when the reward location or environment is changed, LEC neurons largely maintained their firing patterns relative to the new reward, indicating consistent representations of pre-reward and post-reward epochs that are invariant to location and environment and suggesting potential specialized circuitry within LEC.

### LEC pre-reward ramping activity is partly composed of behaviorally linked population state transitions

The pre-reward period, during which the mouse may be anticipating reward, is a particularly interesting reward experience epoch. Both behavioral changes (slowing, Figure 2a) and the firing in pre-reward neurons in LEC (ramping, Figure 4b) precede the reward, which led us to investigate two questions: first, what are the dynamics of population activity of the pre-reward cells, which may give insight into to the internal state of the network; second, does this LEC state relate to the behavior of the animal in a way that might help precisely define the pre-reward epoch.

**Figure 4:**
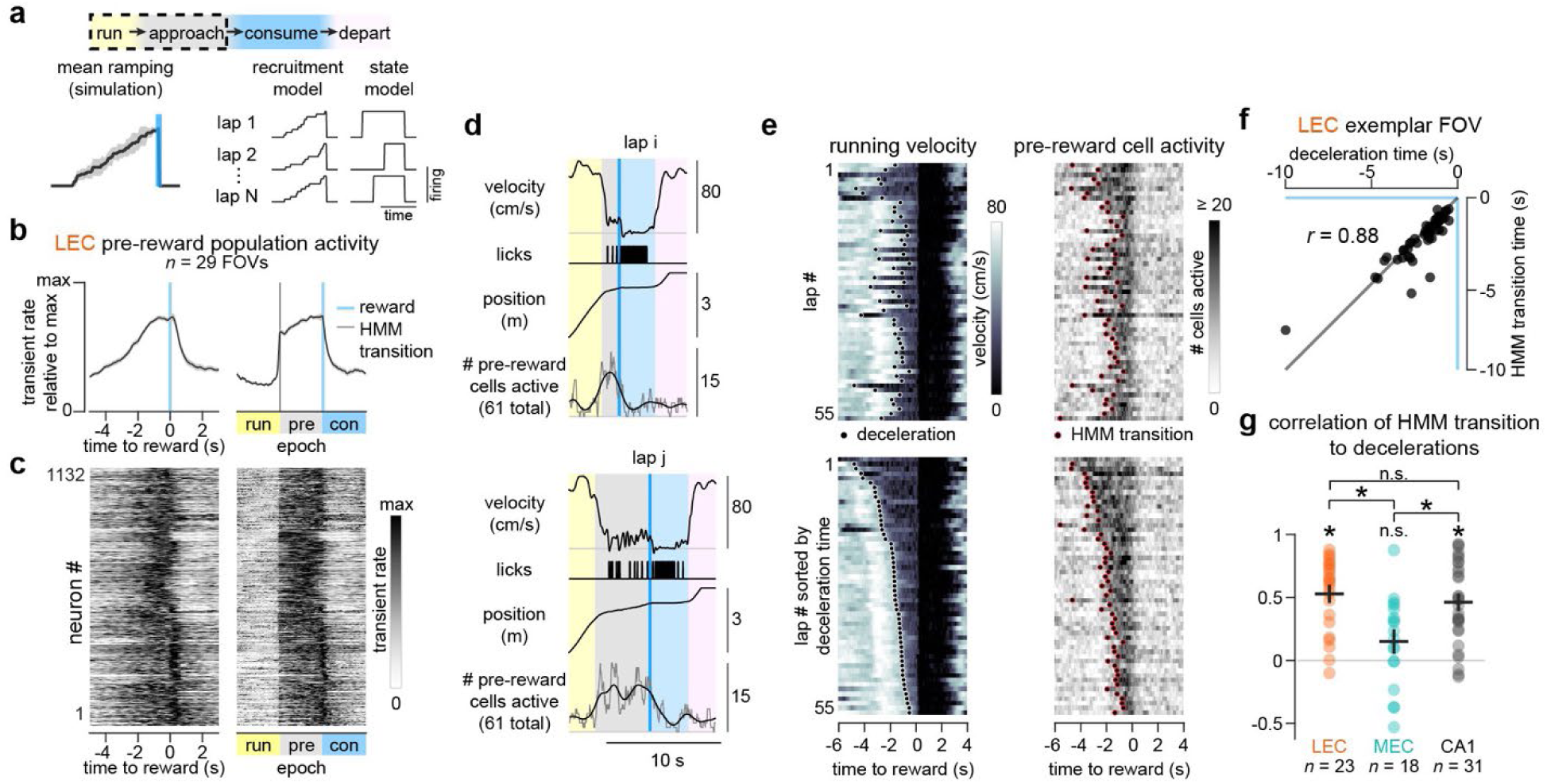
Pre-reward cell activation is partly composed of state changes linked to behavior. a) Pre-reward population activity exhibits a ramp up until the reward delivery. Two potential models that can give rise to mean ramping activity are shown. In a recruitment model, cell activity or the number of active cells increases as the reward location is approached on each individual trial. In a state model, the whole population activates together at some location before reward, but this location is variable across trials. b) Mean activity of LEC pre-reward population as a function of time relative to reward or by epoch, where the three epochs are defined by the start of the traversal, the HMM-detected transition of pre-reward cells to an active state (as in e), and the reward delivery time, and intervals within each epoch are linearly interpolated across time between the transitions. Data is calculated for individual sessions (mean of all pre-reward cells) and normalized to their maximum values; mean ± SEM are shown as dark line with light shading, respectively. c) Activity for individual neurons by time or by epoch. Each row is data for one LEC pre-reward neuron. Data is averaged across laps in a session, normalized to the maximum, and grouped by session. d) Representative sample of behavior and pre-reward activity during two traversals on virtual track. The number of pre-reward cells active (out of 61) is shown as the raw value (gray trace) and after applying a 2 s Gaussian filter (dark trace). e) Running velocity and number of pre-reward cells active on each lap. Detected deceleration and HMM-detected state transition times are shown with black and red dots. Top plots are sorted by lap number for this session. Bottom plots are sorted by deceleration times on each lap. f) Times of deceleration and state changes for exemplar session from e. Each point represents data from one lap. *r* indicates Pearson correlation coefficient. g) Pearson correlation coefficients for each session, calculated as in f, for fields with at least 10 pre-reward cells. Each point represents one imaging session; black cross represents mean ± SEM across sessions. Statistical tests performed between each pair of regions (2-sample t-test) and to zero as well (1-sample t-test).

On average, mean firing in pre-reward LEC neurons ramped up until the reward was delivered (Figure 4b), similar to activity previously observed in other brain regions such as the ventral tegmental area^45^ and the orbitofrontal cortex^46^. We first asked what underlying population dynamics could give rise to this observed mean ramping signal. Various models have been proposed^47–49^. For example, an increasing number of neurons might be recruited or firing in individual neurons could increase as the animal approaches reward, in which case ramping activity would be observed across the population on both individual trials and the trial averages (recruitment model, Figure 4a). Alternatively, the pre-reward population might undergo coherent state changes at different times or positions with respect to reward on each trial, in which case discrete changes in activity, not ramping, would be observed across the population on individual trials and ramping would only be observed in trial averages (state model, Figure 4a). To examine these possibilities further, we quantified the number of pre-reward neurons active in the period leading up to reward on individual trials. We often observed step-like increases in the active number of neurons (Figure 4d-e). Indeed, when we used a hidden Markov model (HMM) to identify such transition times from inactive to active population states (Supplementary Figure 4a) and then plotted the firing rate of the pre-reward neurons aligned to these transition times (with each ‘epoch’ spanning the time between transitions using linear interpolation), we observed a clear step-like increase across the population of individual pre-reward neurons (Figure 4b,c). We further calculated that 44% of the variance in single-lap activity was explained by a step increase. However, some ramping activity was still observed on top of the step-like change (Figure 4b). Thus, part of the increasing ramp in average LEC pre-reward neuron activity could be explained by a discrete switch in firing state occurring at variable times and positions with respect to reward on different traversals, but some contributions from an increasing number of active neurons (or an increase in firing in individual neurons) as the animal approaches reward were also present.

Next, we asked whether the state transitions in LEC related to the behavior of the animal. Interestingly, on individual trials we also observed state changes in mouse behavior: mice switched from a fast running speed to a much slower speed ahead of the reward location and maintained this slow speed until the reward was encountered (Figure 4d and Supplementary Figure 4b-c), with the timing and position of this deceleration varying across trials (Figure 4e). To investigate whether the discrete behavior changes correlated with the LEC population state changes, as suggested by individual example trials (Figure 4d), we examined the relationship between the HMM state transition times with the deceleration times on each trial (Figure 4e). We found these times were highly correlated, both for the example session (Figure 4f) and across all LEC sessions (Figure 4g). A similar effect was observed for CA1 pre-reward cells, but no correlation was observed for MEC pre-reward cells (Figure 4g). Thus, the discrete transitions in population firing state in LEC pre-reward neurons correlates with slowing behavior.

### Location-invariant representation of reward consumption in the LEC

Pre-reward and post-reward periods are separated in time by a period of reward consumption, during which the animal is stationary and consuming water. Signaling in this epoch (during the successful discovery and consumption of a reward) is necessary to forming a representation of the experience of the reward. Since this epoch is typically excluded from analysis of spatial coding properties of cells (which excludes times when velocity is zero), we examined LEC firing relative to reward time, focusing on the period of reward delivery and consumption, in order to investigate how LEC represents this reward consumption epoch. At the time of reward delivery, we observed that LEC population firing increased dramatically and transiently, a signal not prominent in either MEC or CA1 (Figure 5a). Nearly 1 in 8 LEC neurons peaked within the first second after reward delivery. Using a shuffle test for significance of this firing peak, we developed criteria for ‘reward consumption active’ (RCA) neurons^20^ that contain such a peak in firing at reward, or RCA cells. Across all sessions, 13.7% of active LEC neurons (1978 of 14489) qualified as RCA cells (Figure 5b-c), and this proportion was similar between layers II and III of the LEC (Supplementary Figure 2b). The fraction of RCA cells was much lower in the MEC (299 of 9488, or 3.1%), while CA1 was intermediate (816 of 11564, or 7.1%) (Figure 5d).

**Figure 5:**
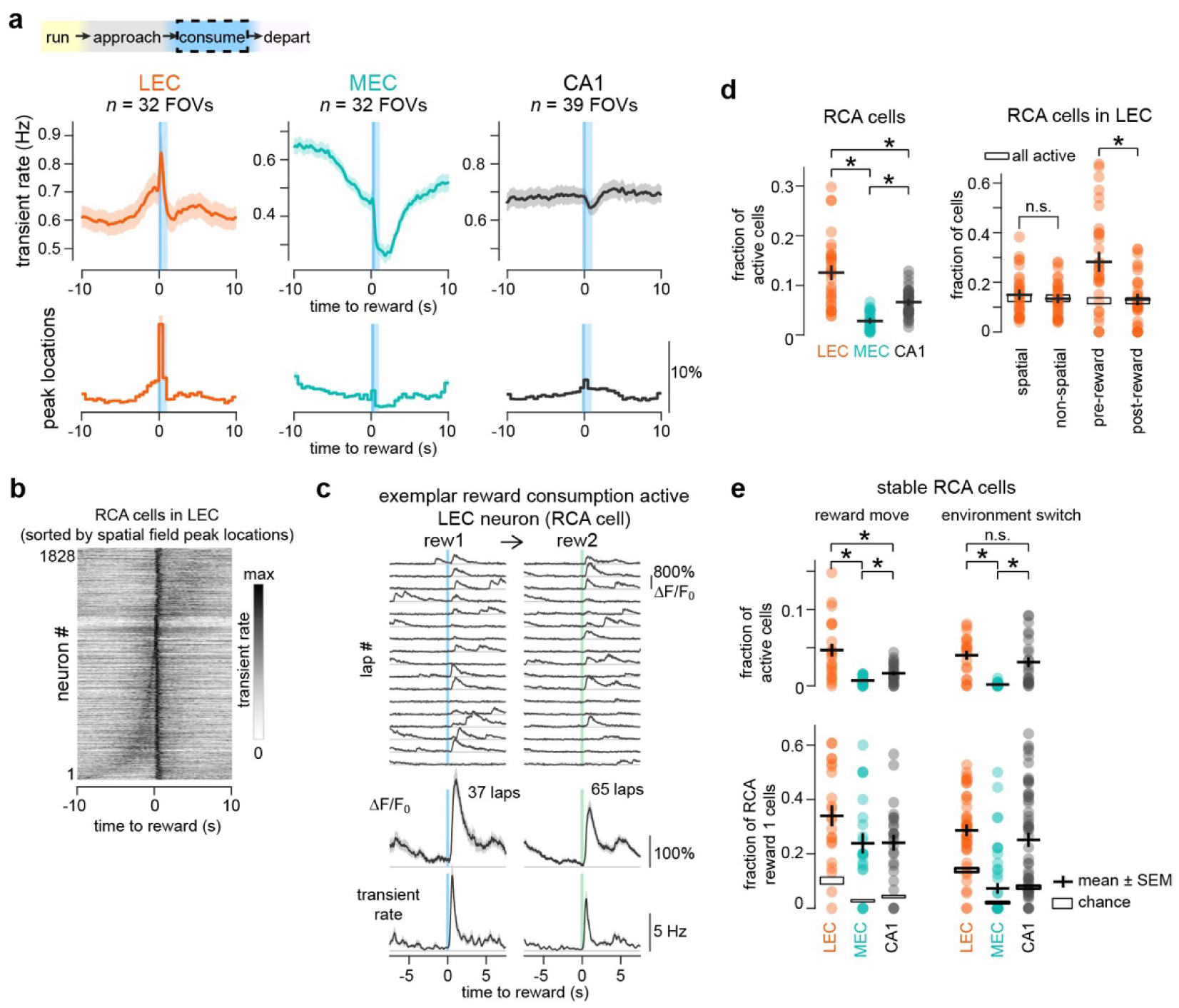
A population of LEC neurons signal reward consumption. a) Mean transient rate relative to the time of reward delivery, shown for LEC, MEC, and CA1 imaging fields averaged by imaging session. Bottom plot is a histogram of the timing of peak firing for individual cells from each session. All active cells are included. Mean ± SEM are shown as dark line with light shading, respectively. Blue bar highlights first second after reward delivery. b) Reward consumption active (RCA) LEC neurons (selection criteria in Methods) from all imaging sessions are sorted by their peak firing location along the track, averaged over all laps for a session. Each neuron (one row) is normalized to its maximum firing rate in this time window. c) Exemplar non-spatial RCA cell in LEC. Fluorescence traces for a subset of laps are shown, with averages of all laps underneath for both fluorescence and inferred transient rate following deconvolution relative to a familiar reward location (rew1, blue) and a new reward location (rew2, green) in the same imaging session. Mean ± SEM are shown as dark line with light shading, respectively. d) Fraction of cells that are RCA cells in each region, further subdivided as a fraction of spatial, non-spatial, pre-reward, and post-reward cells in LEC. Each point represents one imaging session; black cross represents mean ± SEM across sessions. Statistical tests performed between pairs of regions (2-sample t-test) or paired tests within LEC (1-sample t-test). * indicates p < 0.05 and n.s. indicates p > 0.05. e) Fraction of stable RCA cells (reward consumption active for both a familiar and a new reward location), either as a fraction of all active cells or as a fraction of RCA cells for the familiar reward location (rew1). Data is shown for both a reward move in the same environment or for an environment switch. Each point represents one imaging session; black cross represents mean ± SEM across sessions; black rectangle represents mean ± SEM expected by chance. Statistical tests performed between each pair of regions (2-sample t-test). * indicates p < 0.05 and n.s. indicates p > 0.05.

Given that the reward consumption epoch, which determines whether a neuron is an RCA cell, is excluded from the calculation of spatial cells, we next asked whether and how the RCA cell population overlapped with spatial cells in LEC. We found that similar fractions of spatial and non-spatial cells were RCA cells (Figure 5d). The non-spatial ‘pure’ RCA cells, such as the exemplar shown (Figure 5c), were robustly active only during reward consumption. As for spatial cells, RCA cells were more common within the pre-reward cell population of LEC, but they were found within the post-reward population as well (Figure 5d). Importantly, when the reward location was moved, the reward consumption signal occurred at the new reward (Figure 5c,e), with a third of RCA cells in LEC maintaining their firing pattern with respect to reward time, higher than in CA1 and MEC (Figure 5e). When the environment was changed, again RCA cells in LEC maintained their firing pattern with respect to reward time in the new environment (Figure 5e). Therefore, we find that LEC represents the epoch of reward delivery and consumption with a dramatic transient increase in firing that is generated by a population of neurons with a significant level of dedication to providing this signal.

To investigate whether the signals in LEC surrounding reward delivery were due to reward itself or to the experiences surrounding reward, in separate experiments we measured the reward-triggered signal in LEC for randomly delivered rewards outside of the context of virtual reality-based spatial navigation. In these sessions, we still observed a dramatic and transient increase in firing immediately after reward delivery (for 1 s), as seen for RCA cells, but did not observe the increasing ramp in firing leading up to reward delivery nor the later increase in firing 2-4 s after reward delivery (Supplementary Figure 6). Thus, the signal carried by RCA cells appear to be a generalizable reward signal while the pre-reward and post-reward signals are specific to our navigation task.

**Figure 6:**
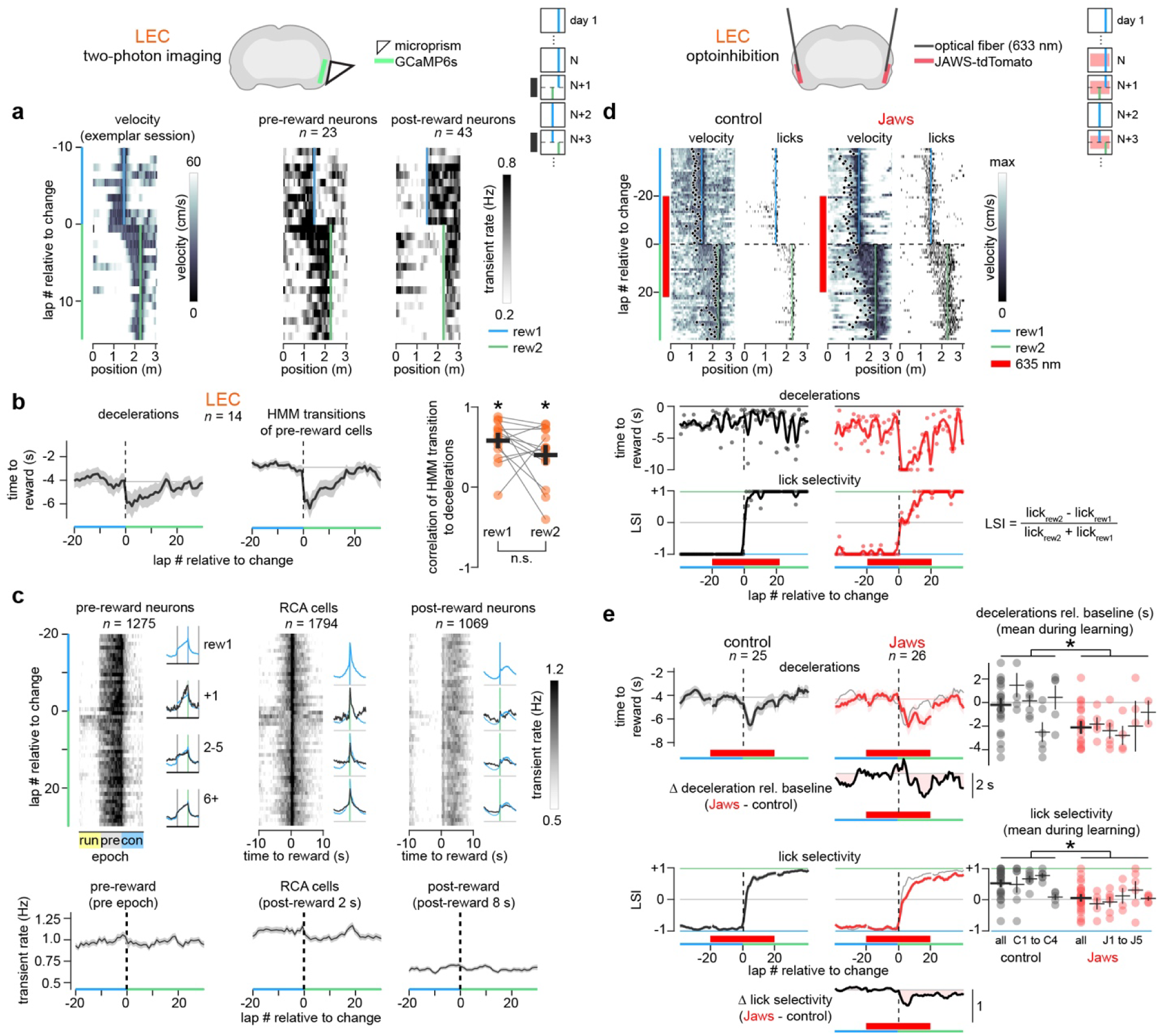
LEC stably represents reward experience during learning while optogenetic inhibition of LEC disrupts learning. a) Mice learned a new reward location over the course of a few laps. In this exemplar session, the reward location is moved later along the track. Velocity along the track, binned at 1 cm intervals, is shown for the final 10 laps with the familiar reward location and the first 15 laps with the new reward location. Pre-reward and post-reward population firing as a function of track position is similarly shown as mice learn the new reward location, averaged over all neurons for that population in this exemplar session. Inset shows imaging paradigm (similar to Figure 3a). b) Line plots show deceleration and HMM transition times during the learning period, averaged across imaging sessions (only sessions with at least 10 pre-reward neurons are included; further, the first session with any reward location move is excluded for each mouse). Mean ± SEM are shown as dark line with light shading, respectively. Pearson correlation coefficients between pre-reward HMM transition and decelerations on each lap, shown for all laps with the familiar reward location (rew1) and the new reward location (rew2). Each point represents one imaging session in LEC; black cross represents mean ± SEM across sessions. Statistical tests performed between rew1 and rew2 conditions (Wilcoxon signed-rank test) and to zero as well (1-sample t-test). * indicates p < 0.05 and n.s. indicates p > 0.05. c) Mean firing for pre-reward neurons, RCA cells, or post-reward neurons. For pre-reward cells, data resampled as a function of behavioral epoch, shown for 20 laps before and 30 laps after reward location change. Insets show firing averaged over subsets of laps (rew1: last 20 laps with familiar reward location; +1: first lap after reward location change; 2-5: next 4 laps; 6+: remaining laps with new reward location). Underneath, mean transient rate for LEC pre-reward cells as a function of lap number, averaged over the pre-reward epoch. For RCA or post-reward cells, mean firing shown as a function of time relative to reward delivery, with mean transient rate averaged over the first 2 s after reward delivery for RCA cells or the remaining time after reward (2 to 10 s) for post-reward cells. d) Velocity and licking behavior for exemplar sessions for control (mCherry injections) and Jaws mice. Trials with optogenetic inhibition by 633 nm light are indicated by red bar (light delivered over entire track traversal for these trials), which is delivered for the final 10-20 laps with the familiar reward location (rew1, blue line) and the first ~20 laps with the new reward location (rew2, green line). Velocity max for the color scale is 30 cm/s for the control and 50 cm/s for Jaws. Underneath, the lap-by-lap measures of deceleration time relative to reward and the lick selectivity index (LSI, Methods) are shown, with dots representing the value for individual laps and the line the smoothed data (5-point rectangular filter). LSI ranges from −1, indicating 100% of the licks are near the familiar reward location, and +1, indicating 100% of the licks are near the new reward location. Inset shows paradigm across days. At least one session was conducted between reward location changes to allow familiarity of that reward location. e) Session averages of deceleration times relative to reward and lick selectivity. For Jaws data, the mean control trace is reproduced using light gray for comparison; moreover, the difference between Jaws and control data is shown underneath, with baseline adjusted for the deceleration data for each session (calculated as the mean deceleration times for laps before reward switch but with 633 nm light on). “Mean during learning” quantifies mean value of deceleration times (on laps 13-20 after reward is moved) relative to baseline (calculated as mean deceleration times on the 10 laps before reward is moved) and mean value of lick selectivity on laps 2-6. Each point represents one imaging session; the first session with any reward location move is excluded for each mouse. Data is shown both for all mice (‘all’) and broken up by individual mice (C1 to C4 are control mice; J1 to J5 are Jaws mice). Black cross represents mean ± SEM across sessions. * indicates p < 0.05, 2-sample t-test. n.s. indicates p > 0.05.

### Trajectory of LEC population firing encodes the reward experience

Inspired by a prior study using a cue-reward association task to investigate layer II fan cells in the LEC^35^, we applied principal component analysis to our population of 9554 active LEC neurons for the 22 imaging sessions performed during a change in reward location and visualized the LEC population firing using first two principal components (PC1 and PC2), which explained 20% and 10% of the variance, respectively (Supplementary Figure 7). The LEC population firing was plotted with respect to reward experience epochs as established in Figure 4. We made two observations. First, in this state space, jumps in the neural activity trajectory occur during transitions between the (previously defined) experience epochs (post-reward/running to pre-reward approach to consumption), which was quantified as the magnitude of the difference between successive points (Supplementary Figure 7b). Second, after the reward was moved, the neural activity was largely unperturbed, following a similar trajectory as for the familiar reward (Supplementary Figure 7a). This observation was quantified by taking the difference between the state space trajectories in the first two principal component dimensions (Supplementary Figure 7c). For reference, we compared this quantity to the same measure applied to MEC and CA1. Unlike LEC, the trajectories in MEC and CA1 differed more after the reward location was moved. Thus, this state space analysis adds further support to our conclusions about LEC coding of reward location but from a new perspective; as a population, the LEC population robustly tracks reward experience epochs.

### Stability of LEC reward representation during learning

Thus far, we have established that three different LEC populations – pre-reward, post-reward, and reward consumption active – largely maintain their same representation around different reward locations. We next investigated the lap-by-lap changes in these representations following a reward location change to determine if the cells quickly and stably encoded the new reward or whether they slowly formed their representations, as seen for CA1 place cells in prior studies^27,50^.

As mice learned a new reward location, they adapted their behavior by slowing down for a longer time and distance before the reward and running slowly until receiving the reward (Figure 6a-b). Over subsequent laps, the mice slowed closer and closer to the new reward zone, with their running behavior converging to the same stable pattern observed around familiar rewards, but relative to the new reward location. While mouse behavior adapted over several laps, throughout this time the pre-reward population and the post-reward population continued to fire before and after the new reward location on each lap (Figure 6a).

In particular, the pre-reward population appeared to evolve during the learning period. Initially, these cells were broadly active across a large portion of the track, which gradually sharpened to be active only in locations just before the reward (Figure 6a). This switch from inactive to active on individual laps, fit with an HMM as in Figure 4, captured this evolution as well (Figure 6b). Moreover, activation of the pre-reward cells was highly correlated to slowing behavior on individual laps for both the familiar and the new reward locations (Figure 6b). Indeed, when reexamining cell firing with respect to the approach epoch, defined by the time from deceleration until reward delivery (as in Figure 4c), the stability of the pre-reward representation across the switch becomes evident (Figure 6c). When mean activity is evaluated for this pre-reward epoch, no detectable change in firing rate across the pre-reward population was observed during learning of the new reward location (Figure 6c).

Surprisingly, both the RCA population and the post-reward population began encoding the reward consumption epoch immediately on the first traversal after the switch (Figure 6c). Even though the receipt of reward was unexpected at the new location, the amplitude of the transient increase in firing in the RCA population did not change across the reward switch laps (Figure 6c). The post-reward population rapidly shifted to fire at the new reward location (Figure 6a), thus encoding the new post-reward epoch from the first traversal after the switch and with no detectable change in firing rate (Figure 6c).

Overall, these results indicate that pre-, post-reward and RCA populations in LEC shift their firing immediately to the new reward location during learning. Given such stable location-invariant reward representations, even while reward location and animal behavior dramatically changed, the firing patterns of these neurons provide reliable experiential information of the epochs surrounding reward.

### Inhibition of LEC disrupts learning of a new reward location

Given the stable representation by LEC of the experiential epochs surrounding reward during learning of a new reward location, we asked whether LEC itself was causally involved in the behavioral changes that occur after reward switch. We expressed the inhibitory opsin Jaws^51^ or a nonfunctional fluorescent marker (mCherry) as a control in LEC and, using 633 nm illumination delivered to chronically implanted fibers in bilateral LEC, inhibited LEC activity in the 10-20 laps prior to moving the reward and through the first 20 laps during learning of the new reward location (Figure 6d, Supplementary Table 3). We found no change in behavior when we inactivated LEC before moving the reward location, as assessed by anticipatory deceleration and licking selectivity (LSI, Methods) at the reward location. When the reward was moved, control mice quickly adapted their behavior to the new reward location, shifting their decelerations and licking within the first ~10 laps (Figure 6d-e), similar to the behavior seen in mice during two-photon imaging of the LEC (Figure 6b). However, mice expressing Jaws took longer to adapt their behavior to the new reward location (Figure 6d-e). Both deceleration behavior and licking selectivity adapted more slowly when compared to control mice. These results indicate that LEC is necessary for the learning of new reward locations but not for already learned reward locations.

## Discussion

The original proposal of a cognitive map^52^ described not just a representation of spatial position but information to guide goal-directed navigation, a dichotomy exemplified by the context-dependence of place cells in the hippocampus^9,10^. In particular, a representation of reward experiences provides information needed to contextualize a spatial code^1,2^. How this reward experience information reaches the hippocampus has been uncertain. Here, by developing a novel imaging approach (Figure 1), we find that the LEC contains cell populations that signal goal approach, the reward consumption period itself, and goal departure; these populations are largely invariant to spatial location or environment (Figures 3 and 5); these populations maintain stability during learning; and inhibition of LEC significantly slows learning of a new reward location (Figure 6). We further show that these signals differ drastically from the MEC, which shows little change around reward and does not represent reward experience epochs in an obvious way.

Previous studies have found a role for LEC in tasks with some similarities to the goal-directed navigation paradigm used here. Ablation of the LEC impairs association of objects with context^53^ and location^54^, LEC neurons signal goal approach in open field navigation^55^, and the LEC is involved in forming associative memories^35^. Our results add to these previous findings, particularly by dissociating reward from object representations. We find that, among other roles, the LEC can provide the “what” information of an experience^56^, which during reward-guided navigation correspond to the epochs relative to the reward. And, consistent with past lesioning studies in the hippocampal formation^57,58^ and the role of LEC in associating objects with location^54,54^, silencing LEC disrupts learning of a change in reward location but does not disrupt behavior around a reward location that is already learned. The LEC, therefore, provides information about the reward experience that the hippocampus could use alongside MEC spatial inputs. By providing these stable representations of reward experience in parallel to spatial information, the entorhinal cortex could provide a computationally efficient and flexible system for learning to associate them^5,7^. A novelty signal could then be provided by neuromodulatory systems such as the locus coeruleus^29,30^ or VTA^59^, which can open a window for plasticity when needed, such as when encountering a new environment.

The reward experience information from our three functional populations of LEC neurons could encode the behavioral actions of the animal, the sensory experiences during that epoch, or some other internal or latent state. For example, the pre-reward population may represent deceleration behavior, visual inputs that predict the reward, or anticipation of the reward itself. Irrespective of which information is represented, the signal is informative to the hippocampus. A further elaboration of this idea is that each functional population may be involved in a more active computation during its respective epoch beyond just signaling the experiential information, and these specializations may have implications for each population’s role in the hippocampal circuit. We briefly discuss here some of the evidence and implications for these specializations in each cell population.

During goal approach, activation of the pre-reward population coincides with the deceleration of the animal (Figure 4). A possibility is that the pre-reward population is only indicating a read-out of running speed, but three properties of this population argue for broader encoding properties. First, these cells are enriched in layer II of the LEC. This layer contains the dopamine-recipient fan cells^35^, which project to dentate gyrus^34^, and pyramidal cells that project to CA3^21^. Thus the circuitry of this population appears to be uniquely positioned to be modulated by direct dopamine release during reward anticipation^45^. Further research may delineate whether fan cells themselves are enriched in the pre-reward population and, if so, what role dopamine plays in modulating firing in this population during goal approach^60^. Second, the goal approach epoch contains rich and interesting behavior, reflected in the slower running speed and variable deceleration times, which may indicate the need for gathering more information for optimal foraging^61,62^. During this period, the LEC population firing increases, opposite of what we observed in the MEC. Finally, despite the increase in firing, the LEC population decorrelates during goal approach, consistent with increased information coding (Supplementary Figure 5)^63,64^. Together, this evidence points to the goal approach epoch and especially the LEC pre-reward population as playing an important role for the hippocampus and other downstream regions in forming memories of the experience and improving predictions that could guide future behavior^7,65^. This pre-reward population shares many qualities with cue-active cells previously identified in the LEC using a different task^35^. These cells – identified among fan cells in layer II of the LEC – fired in the period between cue onset and reward delivery, consistent with our results. Thus, this population may be encoding expectation or value, which may be generated in cooperation with signals from dopaminergic cells^66^.

During reward consumption, firing in LEC increases transiently and dramatically (Figure 5 and Supplementary Figure 6). This signal is observed in a population of neurons we have termed RCA cells. These cells are widespread: they are found in both layer II and layer III of the LEC, in both spatial and non-spatial cells, and in both pre-reward and post-reward populations. While CA1 on average did not exhibit a similar increase, perhaps reflecting the balance of increased drive from LEC with decreased drive from MEC (Figure 5a), a fraction of CA1 neurons qualified as RCA cells, which may be neurons that receive stronger inputs from LEC than from MEC^67^. Thus a large burst of excitation may be sent from the LEC to the hippocampus during the initial moments of reward consumption and could drive behavioral timescale synaptic plasticity^68^ or dendritic spikes^69^ in CA1 pyramidal neurons and in turn drive an overrepresentation of reward locations^27^.

During goal departure, post-reward cells in LEC became active. Interestingly, MEC and CA1 were also enriched in the number of post-reward cells (Figure 2). Along with signaling the goal departure experience itself, neurons active in the post-reward epoch might be coordinated across all three regions to provide a trace signal, which could be used to associate post-reward locations with the reward^70,71^. An alternative interpretation is that the post-reward signal is due to reward consumption or satiation and has no special role in spatial navigation; however, the absence of a prominent post-reward signal for randomly delivered rewards outside of a task structure (Supplementary Figure 6) supports the notion that post-reward firing is specific to encoding information that relates to predicting the goal location during future behavior.

Our findings revisit the discussion about the dueling identities of MEC and LEC and how to frame the distinction (or the similarities). These differences have been cast in terms of “where” versus “what”, spatial versus nonspatial, self and non-self, and more^56,72,73^. Recent work found support for an allocentric versus egocentric distinction^55^. During open field navigation, neurons in LEC were found to encode egocentric coordinates relative to items in the environment, which contrasted with the allocentric encoding of neurons in MEC. A parsimonious explanation of our findings, particularly the populations of pre-reward, RCA, and post-reward cells, is that these cells encode egocentric coordinates, but rather than representing position relative to physical objects they represent position relative to a more abstract object, which is the reward itself (or, more concretely, when and where the reward experience will occur). Indeed, the past study identified goal-related egocentric tuning. We propose a subtle yet important distinction to this interpretation. Our finding that pre-reward populations are activated in a behaviorally-dependent manner argues that the proper coordinate system to consider is not a physical distance metric (such as reward vector coding) but instead a more abstract or internal behavioral space that we describe broadly as experiential epoch coding, consistent with a past proposal that LEC encodes the “content of an experience”^56^. Interestingly, previous work that had found a prominent trajectory encoding signal in rat entorhinal cortex during reward navigation^12^ was likely sampling LEC, not MEC, consistent with our findings and further distinguishing the role of these two regions.

Our results provide an important complement to recent papers studying the role of LEC in reward-based tasks. First, during a task requiring rats to hold a lever before receiving reward, multiple reward-related signals were observed in the LEC and CA1^74^, including “hold-type” cue-active cells that may be similar to the pre-reward cells we observed during reward guided navigation. In another study, during an odor cue-reward association task, cue-active cells were found in fan cells of the LEC (which exclusively reside in layer II)^35^, similar to our finding of an enrichment of pre-reward cells in layer II of the LEC. Thus, despite the differences in our tasks and our recording techniques, the same cells in LEC may generalize to perform a similar function. Beyond these parallels, our findings also provide novel insights. By using a spatial navigation task, we were able to describe a number of new categorizations that go beyond what was described in prior research. While our pre-reward cells may parallel the cue-active cells mentioned above, we describe a behaviorally-linked state transition in pre-reward firing, and variability in the timing of state transitions contributes to the observed ramping activity (Figure 4). This result may be important not just for reward navigation signals in the hippocampus but may be of interest to studies of ramping signals in the dopamine system^47,75^. Further, analogues of our reward consumption active cells and post-reward (goal departure) cells have not been previously described. This latter group of goal departure cells may be unique to spatial navigation tasks.

In another recent study, axons from LEC and MEC were imaged in CA1 during a reward navigation task in virtual reality^76^, similar to the task used here. They found that the CA1-projecting axons from layer III of the LEC represented reward but also strongly coded spatial position as well. Our approach complements these results as our imaging method allows us to sample both CA1-projecting layer III cells and layer II stellate and pyramidal cells that project to the dentate gyrus and CA3, although our method does not allow us to confirm projection targets of individual cells. Our results indicate layer III of LEC does indeed carry more spatial information than layer II (Supplementary Figure 2b), thus explaining why a relatively high degree of spatial information was reported in CA1-projecting LEC axons^76^. Together our methods provide a more comprehensive picture for how information from LEC is reaching the hippocampus during reward navigation behaviors, with layer II providing the strongest pre-reward information that is likely routed to DG and CA3 while layer III is providing stronger spatial information that is routed to CA1. Both layers contribute reward consumption and post-reward information.

Our results help bridge a gap between models of goal-directed navigation and known properties of the entorhinal-hippocampal circuit. For example, a critical component of the successor representation model of the hippocampus^7^ is the presence of a source of reward prediction information^1^. The presence of pre-reward, reward consumption, and post-reward cell populations strongly implicates the LEC as the source of this reward information. Further, the preservation of these signals after moving the reward location or changing the environment completely lends support for dedicated functional modules within the LEC. Such organization has implications for computational models of the entorhinal cortex^77^.

Finally, the LEC is known to be involved in many tasks and behaviors beyond what was studied here, such as in olfaction^34^ and timing on the order of minutes^33^. Indeed, the spatial and reward-encoding populations we identified represent only a third of neurons active during our spatial navigation task. Thus, it is yet to be determined precisely how these diverse roles are supported by the same brain region or what the remaining two-thirds of neurons are encoding. One possibility is that different subcircuits within the LEC each specialize in encoding a particular type of information or experience but, without behavioral measures of these other aspects of experience, it was not possible here to determine their coding. Instead, we first identified a subset of neurons with “spatial” information. In actuality these neurons encoded the reward experience, which changed with spatial position and thus our analysis could identify their firing properties. Our findings also provide support for the hypothesis of functional specialization of LEC subcircuits since many pre-, post-reward and RCA cells maintained their encoding across reward and environment switches; thus, perhaps they are recruited from largely non-overlapping pools of LEC neurons, each with its own specialized circuitry and dedicated function. Unraveling the precise nature of the LEC’s role across a variety of tasks and behaviors will be made possible by combining large-scale population imaging with an ever-expanding suite of genetically-targeted optical tools.

## Methods

All animal procedures were approved by the Northwestern University Institutional Animal Care and Use Committee. For imaging of the medial and lateral entorhinal cortices (MEC and LEC, respectively), mice expressing GCaMP6s were generated by crossing tetO-GCaMP6s mice (JAX No. 024742) with Camk2a-tTA mice (JAX No. 007004). For imaging of CA1 or optogenetic inhibition of LEC, wild type offspring of these crossings were injected with adeno-associated virus (AAV) as detailed below.

### Surgery

Mice were anesthetized with isoflurane (4% for induction, 1-2% for maintenance in 0.5 L/min O_2_) while temperature was maintained by a heating pad at 37°C as measured by rectal probe. Ointment was applied to the eyes. Dexamethasone (5 mg/kg, i.m.) was given for inflammation, buprenorphine-SR-LAB (1 mg/kg, s.c.) for pain, and normal saline (0.5-1.0 mL, i.p.) for dehydration. Details for the surgeries to implant prisms to access the MEC and the LEC, to inject virus and cannulate to access CA1, or to inject virus and implant fibers to optically inhibit LEC are provided in the next sections. In all cases, after implantation, a titanium headplate was attached to the skull with dental cement (Metabond, Parkell). Mice were monitored closely for 24 hours and given 3-5 days to recover before water restriction and behavioral training were begun.

### Lateral entorhinal cortex surgery

As the LEC lies on the ventrolateral surface of cortex and is surrounded by soft tissue, bony protrusions, rich vasculature, and the pinna, a lateral optical approach is difficult and complicates concurrent head-fixed behavior. Prior work utilized a lateral approach in anesthetized animals and achieved two-photon imaging field sizes of 200 µm and up to 50 neurons at a time^1^. Thus, we strove to develop an imaging approach compatible with behavior that could yield field sizes of at least 500 µm and over 500 neurons at a time. We devised an overhead imaging approach using a microprism to provide optical access to the LEC^2–4^. A 3 mm craniotomy was made over the right lateral surface of the skull and positioned so that the posterior edge aligns with the ventral portion of the transverse sinus and the anterior edge with the insertion of the zygomatic protrusion from the squamosal bone, centered at ~3.5 mm caudal to Bregma. The dorsal edge was 1-2 mm dorsal of the rhinal fissure and the ventral edge was extended as far ventral as possible without incurring large amounts of bleeding or damage to soft tissue structures. Once the brain was exposed, any soft tissue overlying dura was removed. A 3 mm round No. 0 coverslip (CS-3R-0, Warner Instruments) was lowered and held in place with a 1.0 mm diameter pipette (Q100-70-7.5, Sutter Instrument) positioned by a micromanipulator while the top and side edges of the coverslip were cemented (Metabond, Parkell). Then a small dab of UV-cured adhesive (NOA81, Norland) was placed on the outer surface of the coverslip and a 2.0 mm microprism (MPCH-2.0, Tower Optical) placed against the coverslip and positioned as far ventral as possible before UV-curing the adhesive (CS20K2, Thorlabs), thus fixing the microprism in place. Dental cement was then used to fill in the remaining gaps while leaving the dorsal face of the prism clear for optical access.

The location of the window over the LEC was confirmed in three ways (Supplementary Figure 1). First, expression of GCaMP6s is enriched in the entorhinal cortex in the tetO-GCaMP6s x CaMKIIa-tTA transgenic mouse line. We confirmed this overlap through retrograde labeling of CA1-projecting neurons. Labeled neurons in layer III of the LEC and the MEC coincided with increased brightness of GCaMP6s fluorescence in the same regions as visualized in histological slices. Second, this expression and the location of the LEC is found ventral to the rhinal vein. As this is a prominent structure seen during surgical implantation of the window and during subsequent in vivo imaging, it provided a reliable landmark for locating the LEC and, under epifluorescence imaging, matched the area of increased GCaMP6s fluorescence. Finally, the drastic difference in the location and angles of our MEC and LEC prisms (they are perpendicular to each other, with the MEC prism facing anterior and the LEC prism facing medial) ensures that they are targeting different regions of cortex, and this was confirmed by the presence of a lamina dissecans between layers II and III of cortex in only the lateralized and not the posteromedial portions of entorhinal cortex^5^.

### Medial entorhinal cortex surgery

This surgery is a modified version of the MEC prism implant surgery previously described^3^. A 2-3 mm craniotomy was performed over the right cerebellum with the anterior edge positioned along the transverse sinus, just posterior of the lambdoid suture and the medial edge 2 mm lateral of midline. The craniotomy was extended posteriorly and laterally to where the skull begins its ventral descent. To make drilling easier, the mouse was rotated to bring the edge of the craniotomy in plane (counterclockwise roll of 20-30 degrees and downward pitch of 5-10 degrees). Next, a 2 mm incision was made in the dura over cerebellum along the posterior edge of the transverse sinus and the flap of cerebellar dura reflected posteriorly away from the sinus. A portion of cerebellum was then suctioned until the caudal surface of the cortex was clearly visible and expanded to yield a ~2 mm opening. A 45-degree 1.5 or 2.0 mm microprism (MPCH-1.5 or MPCH-2.0, Tower Optical) was mounted onto a custom stainless-steel mount with UV-cured adhesive (NOA81, Norland) and this microprism assembly was wedged with the front surface of the prism abutting the MEC and the back surface against the caudal portion of the skull. Once inserted, the prism assembly could be gently adjusted to achieve maximal exposure of the MEC, angled to match the natural surface of the MEC and to maximize optical clearance for imaging (typically at a clockwise roll of 10 degrees and an upward pitch of 10-15 degrees). Dental cement was then applied around the prism and surrounding skull to hold the prism in place; gentle anterior pressure was applied against the posterior edge of the prism assembly to provide some mechanical stability against the MEC.

### CA1 cannulation

A small craniotomy (~0.5 mm) was performed at 2.3 mm caudal and 1.8 mm lateral (right hemisphere) relative to Bregma. The virus, pAAV.Syn.GCaMP6f.WPRE.SV40 (Addgene catalog #100837-AAV1, diluted ~10x from 2e13 GC/ml stock into phosphate buffered solution (PBS)), was injected by a beveled glass micropipette at a depth of 1.3 mm beneath dura. Typically, two injections, each 60 nL, were performed at spots ~500 µm apart within the same craniotomy. Next, typically in a separate surgery 2-4 days later, a stainless steel cannula with a glued (NOA81, Norland) 2.5 mm No. 1 glass coverslip (Potomac Photonics) was implanted above the hippocampus.

### Retrograde labeling

CA1-projecting neurons were labeled by injection of rAAV2-Retro-CAG-TdTomato (Janelia, diluted 20x from 1.8e12 GC/ml stock into PBS) into the right CA1 using the same procedures for injection of virus as detailed above (“CA1 cannulation”). For in vivo imaging, an LEC prism was implanted on the same day using the procedures as above for “Lateral entorhinal cortex surgery,” and three weeks later two-photon imaging was performed. For histology, injection of virus into CA1 was performed in a transgenic mouse expressing GCaMP6s. After three weeks, the mouse was anesthetized with 5% isoflurane and perfused with 4% paraformaldehyde (PFA). After leaving the extracted brain in PFA at 4°C overnight, it was moved to 30% sucrose in PBS for a few days. Using a freezing microtome, 50 µm horizontal slices were cut and placed on slides. Images were taken with a slide scanner microscope (VS120, Olympus).

### Implantation of optical fibers for optogenetic inhibition of LEC

Inhibition of LEC was achieved by bilateral expression of the inhibitory opsin Jaws^6^ followed by implantation of a pair of tapered optical fibers^7^. First, a craniotomy was performed over right LEC as detailed in “Lateral entorhinal cortex surgery.” The head was rotated 40 degrees (counterclockwise roll), allowing direct access to LEC using a pipette from above. The pipette was positioned ~3.5 mm caudal to Bregma and punctured dura just ventral to the rhinal vein. The pipette was then advanced ~0.5 mm for the first injection and another 0.5 mm for a second injection. Either AAV2/8-hSYN-JAWS-tdTomato-ER2 (Neurophotonics, diluted 2.5x from 9.8e2 GC/ml stock into PBS) or AAV8-hSyn-mCherry (Addgene catalog #114472, diluted ~6x from 2.6e13 GC/ml stock into PBS) was injected for a set of Jaws mice and a set of control mice, respectively. The pipette was withdrawn after 5 minutes. After another 10-15 minutes, the head rotation was reduced to 15 degrees (counterclockwise roll) and a fiber cannulae (Lambda-B Fiber: 0.39 NA, 200 µm with a slotted 1.25 mm ceramic ferrule, 1.5 mm active length plus 1.5 mm tapered implant length; Optogenix) was inserted. The fiber was positioned at ~3.5 mm caudal to Bregma and punctured dura 1.5 mm dorsal to the rhinal vein before being advanced 3 mm and then secured in place with dental cement. The procedure was then repeated for the left LEC. After experiments were completed, horizontal brain slices were taken as detailed above under “Retrograde labeling.”

### Behavior

Water restricted mice received 1.0 mL of water per day. Weights and health were monitored daily. Training and behavior were performed in virtual reality (ViRMEn^8^) and custom-written code (MATLAB, Mathworks) was used to control and synchronize signals with two-photon imaging using a data acquisition card (National Instruments). Head-fixed mice running on a one-dimensional treadmill moved through a virtual reality environment displayed on a set of 5 monitors covering a 225-degree field-of-view (horizontal axis)^3^. Water rewards of 4 µL were delivered at a fixed location on the virtual track. Water volume was controlled by a solenoid that was calibrated based on open duration; typically, a 20 msec opening delivered a 4 µL volume. Once the end of the track was reached, a 4 s “time-out” period was included where the mice were kept at the end of the virtual track before returning to the start of the virtual track. Licks were monitored by a capacitive sensor attached to the lick spout. During imaging experiments, we also monitored the face of the mouse using a camera (Zelux CS165MU1, Thorlabs) synced to the two-photon microscope frame times.

Imaging sessions were performed once the behavior reached a satisfactory level, judged by a) number of laps per minute (> 2 laps per minute in a 40-minute session) and b) anticipation of reward (deceleration and licking before reaching the reward location, judged by examining reward-triggered average of behavior traces). This point was usually reached after 1 to 2 weeks of training (5-10 sessions). On the first 1-3 days of imaging, the reward location was kept fixed (at 2.3 m). Then, in the middle of an imaging session, the reward was moved, and this new position was used for the remainder of the session and the next session as well (Figure 2a). For the next couple weeks, we interleaved more reward location moves between three possible positions: 0.7 m, 1.5, and 2.3 m. Where appropriate, we pooled data from the three reward positions (Supplementary Table 2). We considered the possibility that the direction the reward location was moved (either earlier or later along the track relative to the familiar reward) may influence our results. We repeated the analysis for Figure 3 and Figure 6 for LEC imaging sessions stratified by which direction the reward was moved (earlier: n = 15 FOVs; later: n = 7 FOVs). We did not observe any significant differences. Thus, we pooled data for both directions of reward changes. For environment switch experiments, we began with the same environment used for reward location changes as above (env1), but then moved through the following sequence in a single session: env1-1.5m, env1-2.3 m, env2, env3, env4, and then back to env1-1.5m, with roughly 20 laps (5-10 minutes) in each environment (Supplementary Figure 3c).

To assess the response in LEC to reward delivery and consumption outside of the context of virtual navigation, water restricted naïve mice (*n* = 3) were habituated to head fixation and provided randomly delivered rewards while imaging was performed of the LEC (Supplementary Figure 6). During these sessions, the display monitors were off. After these imaging sessions, one of the mice was trained on the virtual reality navigation task as described above. Imaging was repeated once behavior reached the criteria detailed above.

### Imaging

Widefield images were taken with a custom-built fluorescence microscope with a GFP filter cube through a 2X objective (TL2X-SAP, Thorlabs) and captured with a scientific CMOS camera (Prime BSI Express, Teledyne Photometrics).

Two-photon imaging was performed using a customized microscope with a resonant scanning module (Sutter Instruments) and a 20X objective (LUCPlanFL N, Olympus). Excitation was provided by a mode-locked Ti:Sapphire laser tuned to 920 nm (Chameleon Ultra II, Coherent) with roughly 100 mW of average power coming out of the objective. Emission light was split by a 560 longpass dichroic (FF560-Di01, Semrock) and filtered into red (FF01-620/52, Semrock) and green (FF01-510/84, Semrock) channels before being detected by a pair of GaAsP PMTs (H10770PA-40, Hamamatsu Photonics). Imaging was controlled by ScanImage software (Vidrio). A frame sync signal was sent to the data acquisition card on the virtual reality computer.

Time series movies ranging from 32,000 to 60,000 frames per imaging session were collected at 29.8 Hz. At full magnification, the field size was 700 x 700 µm with 512 x 512 pixels, with most imaging performed at magnifications of 1.0 to 1.5x (maximum of 2.2x). At the start and end of each imaging session, the depth of the imaging field from the surface (dura for entorhinal cortex imaging or the axonal layer for CA1 imaging) was estimated by focusing up using a calibrated micromanipulator to move the microscope relative to the mouse (MP-285, Sutter Instrument). In LEC, we could distinguish layer II and layer III as a cell-free zone separates them, providing a convenient method to differentiate the layers^5^ (Supplementary Figure 1). Such a clear distinction between layers was not present in MEC; we used a threshold of 150 µm below dura to distinguish nominal layer II and nominal layer III. To block stray light from the VR monitors from contaminating the detected emission signal, a light blocking cylinder formed from electrical tape was placed between the headbar and the objective.

### Optical inhibition of LEC during behavior

Once implanted mice learned the task (as described under “Behavior”), optical inhibition sessions were performed. A 633 nm fiber-pigtailed laser diode (LP633-SF50, Thorlabs) was split 50:50 using a fiber optic coupler (TW630R5F1, Thorlabs) and coupled to the implanted fiber cannulae using a pair of patch cables (M83L01, Thorlabs). The laser diode driver (LDC205C, Thorlabs) was used alongside an LD/TEC mount (LDM9LP, Thorlabs) driven by a temperature controller (TED200C, Thorlabs). Driver current was set to produce 6-8 mW of total power coming into each implanted fiber cannulae (typically 130 mA of total current from the laser driver). Power was switched on and off using TTL control of the driver.

The protocol was as follows. A “reward move” session was run as described under “Behavior” but with the inclusion of inhibition laps. On an inhibition lap, the laser light turned on once the mouse reached 30 cm along the track and stayed on until a position of 270 cm. These positions were chosen so that they would encompass all possible reward positions (70, 150, and 230 cm). In a given session, the first 20-30 laps were run with no light with the reward in the familiar position. Then, at least 10 inhibition laps were run (min: 10, max: 25) with the reward still in the familiar position. Next, ~20 more inhibition laps were run (min: 17, max: 23), but now with the reward in the new position. Finally, at least another 20 laps were run with no light and the reward in the new position.

Once mice learned the first reward position (230 cm), we performed an acclimation session with optical inhibition for ~20 laps but with no reward move. The next day, the reward was moved from 230 cm to 150 cm with inhibition laps as described above. Then, we interleaved days with no reward move (and no inhibition) with days with reward moves (and inhibition), thus giving mice an opportunity to adjust to each reward location before the next move.

### Image processing

To extract inferred firing of individual cells from two-photon movies, we performed the following steps: registration to correct for motion artifacts, segmentation to identify cells, extraction to calculate fluorescence signals for each cell (and associated neuropil signals), and integrated iterative inference to estimate an interpretable and decomposed version of cell activity. First, two-photon time-series movies were motion corrected using rigid registration^9^. A target image was found from repeated rounds of registration for a subset of frames (typically 4000). The full movie was then registered to that target image. Cells were then segmented and raw fluorescence traces extracted using a customized implementation of Suite2p^10^ in MATLAB. Movies were downsampled 20x in time (every 20 frames averaged). The first 200 spatial components identified by singular value decomposition of the downsampled movie were then passed into Suite2p (typical parameters: cell diameter = 10 µm, neuropil ratio = 12x or 120 µm). Next, cells were semi-manually curated based on a number of properties, such as size, shape, and brightness. A brightness-over-time signal was produced for each cell mask along with the corresponding neuropil signal surrounding it.

Next, to decompose the activity signal into interpretable units, we used an integrated iterative algorithm we developed to recover an estimate of r, the ratio of neuropil contamination into the cell signal; F0, the baseline fluorescence of the cell; and S, the firing rate of the cell. F0 is the linear summation of a set of basis functions, chosen to be slowly varying sinusoids and exponentials. S is estimated from deconvolution^11^; thus, we assume ΔF/F_0_ is a convolution of S with a kernel. We chose the kernel to be a two-exponential function with ‘on’ and ‘off’ time constants taken from published data^12,13^ for GCaMP6s and GCaMP6f. S is not an exact measure but does correlate with [Ca^2+^] and thus the number of action potentials fired in a given time window^14^. Thus we scaled S by a number based on the transient dynamics to a number that is meant to approximate the number of action potentials (120x for GCaMP6s and 80x for GCaMP6f, chosen so that the mean value of S matches the measured number of action potentials that generate the same mean ΔF/F_0_ as observed for simultaneous imaging and cell-attached recordings of neurons in vivo^12,13^). The units for S are meant for convenience since the true signals are nonlinear and the dynamics can vary from cell to cell, especially when the sensor is virally expressed^15^. For analysis, S was then smoothed with a gaussian filter (standard deviation of 5 bins or 170 ms).

### Analysis

Analysis was performed in four different reference frames, calculated lap-by-lap and peak locations are defined by the peak in the lap-averages in a given reference frame:

1. Absolute position: calculated by taking frames with positive velocity (at least 10 cm/s) and excluding the first 2 s after reward delivery (if within 10 cm of reward location) and then binning inferred firing S by position into 1 cm bins.
2. Relative position: calculated similar to absolute position but for the 100 cm before and after the reward location, also in 1 cm bins.
3. Relative time: no frames excluded, for the 10 s before and the 10 s after the reward time, binned at 30 Hz. While mice could run the 3.1 m track in under 10 s, they typically took ~20 s (across 2888 laps in our LEC dataset, the mean lap time was 18.7 s with only 1 lap faster than 10 s). This discrepancy is because a) the mice peaked at 50 cm/s or more but ran at a lower speed along a significant portion of the track, b) the mice spent some time (typically 2-4 s) stopped to consume the reward, and c) the end of the track included a “time-out” period of 4 seconds to help separate each lap as a distinct trial. Ultimately, any choice of window is a compromise as it depends on the exact behavior of each mouse. We chose 10 s as a more generous window so that we could ensure we captured most of the dynamics in activity before and after reward delivery.
4. Epochs: formed by defining three events for each lap and linearly interpolating in time 60 bins for each epoch. The three events are start of running, deceleration or HMM transition time, and reward time. These three events were found automatically for each lap by examining the recorded behavior. Start of running after reward was found by examining the position of the animal in the virtual world and finding when the animal first passed 15 cm beyond the reward position. Next, deceleration time (or HMM transition time for the pre-reward population) was found as defined below (under “Deceleration criteria” and “HMM transition criteria”; which measure was used is indicated as appropriate in the text). Finally, reward delivery time was recorded by the virtual reality program and defined the third event. For each inter-event period, we then linearly interpolated the time span between events into 60 bins using the interp1 function in MATLAB.

Criteria for categorizing cells are as follows:

*Active cells*: mean ΔF/F_0_ > 0.1. The fluorescence signal was averaged over all laps and time points for the imaging session.

*Spatial cells*: spatial information >0.3 bits/event and significance on 98% or more of shuffles. Spatial information^16^ was calculated by taking frames with positive velocity (at least 10 cm/s) and excluding the first 2 s after reward delivery (if within 10 cm of reward location) and then binning inferred firing S by position into 60 bins (5 cm). For the shuffled data, firing on individual laps was randomly permuted and the spatial information calculated for the shuffled data. This procedure was repeated 100 times. The spatial information of the cell was considered significant if at most 2 shuffles returned a higher spatial information score.

*Reward cells*: spatial cells with a peak within 40 cm of the reward location (pre-reward: 40 cm before reward and up to and inclusive of reward location; post-reward: after reward and up to and inclusive of 40 cm after reward location).

*Reward consumption active (RCA) cells*: peaks between 0 and 1 second after reward (inclusive), 98% or above on shuffle test for first second after reward (considered significant if at most 2 shuffles out of 100 returned a higher mean firing rate in a 1 second bin than the actual mean firing rate between 0 and 1 s.

Other measures used in the manuscript are defined here:

*Reward clustering ratio*: for spatial cells, the number of cells with peak locations in the zone of interest (reward, pre-reward, or post-reward) was divided by the number of cells with peak locations anywhere along the track, normalized by the expected fraction if cells were uniformly distributed. For example, for a 40 cm pre-reward zone along a 310 cm track, if 20 cells out of 100 have peaks in the pre-reward zone, then the ratio is 20/100 (actual) divided by 40/310 (expected), which yields a pre-reward clustering ratio of 1.55.

*Tuning width*: length of track over which lap-averaged cell firing is greater than 30% of the max.

*Transient duration*: the autocorrelation for ΔF/F_0_ was calculated after mean subtraction and smoothing with a 9-point rectangular filter. The half-height half-width was then found as a proxy measure of transient duration. This measure is of course influenced not just by the kinetics of the Ca^2+^ indicator itself but also by the concentration of the indicator in the cell^17^, endogenous buffers and other properties of each cell, and the autocorrelation in time of the cell’s actual action potential train.

*Deceleration criteria*: 98^th^ percentile of velocity (after smoothing with a 31-point rectangular filter, ~350 ms) was used as the ‘peak’ velocity, vpeak. Vth = vpeak*0.8-5, so if peak velocity was 50 cm/s, we used 35 cm/s as the threshold. We then detected when the smoothed velocity (11-point rectangular filter, ~120 ms) last was above this threshold in a time window from 10 s to 0.5 s prior to reward.

*HMM transition criteria*: for imaging sessions with at least 5 pre-reward cells, we calculated transitions from inactive to active in the 10 s period leading up to reward using a hidden Markov model. The sequence was the number of cells that passed a firing threshold in each time bin (>1.8 events/s). The model assumed only two states, inactive and active, with each lap beginning in the inactive state and transitioning to an absorbing active state. This transition matrix and the probability distributions for number of active cells in each state was estimated using the Baum-Welch algorithm (hmmtrain, MATLAB). Then, for each lap, this model was used to estimate the posterior state probabilities (hmmdecode, MATLAB) and hence the time of transition from inactive to active. To calculate variance explained, we modeled the 10 s pre-reward period for each lap as a step function using the HMM to set the transition point of the step. Then we calculated one minus the variance of the difference between the data and the step function divided by the variance of the data itself: 1 – var(data-step)/var(data).

*Velocity HMM criteria*: we also detected decelerations using a hidden Markov model. Velocity was rounded up to the nearest integer value in cm/s. Similar to the model used above for pre-reward cells, we used two states with a single transition to the absorbing state. In this case, the first state was a high velocity state and the second absorbing state was a lower velocity state. We compared the results of this method to the first method (‘Deceleration criteria’) and also re-ran our analysis comparing HMM-detected decelerations to HMM-detected changes in pre-reward and found both methods produced similar results (Supplementary Figure 4c).

*Population correlation*: cell-cell correlations were calculated at a given position (or time point relative to reward) across laps. For an imaging session, the mean population correlation was then taken as the average cell-cell correlation across all cell pairs (autocorrelations excluded). Positions were binned in 20 cm intervals and time in 1 s intervals.

*Lick selectivity index*: the ratio of licking at the familiar (rew1) or new (rew2) reward locations. L1 is the licking in a zone around rew1, stretching from 30 cm before to 10 cm after the reward location, and excluding any stationary periods. L2 is the licking in a zone around rew2. The lick selectivity index is then calculated as (L2-L1)/(L2-L1) for each lap. A value of −1 indicates licking at rew1 and not rew2 (and the opposite for a value of +1), while a value of 0 indicates equal amounts of licking at both locations. If no licking was detected at either reward location on a given lap, the index was undefined and that particular lap is excluded when calculating averages across sessions.

Bayesian decoding^18^ was performed with respect to absolute position for datasets with reward located at 2.3 m (Supplementary Figure 2c). Data was binned every 10 cm and training and test datasets were formed from odd and even laps, respectively.

## Acknowledgements

We thank Peter Dayan, Jim Heys, Jack Mellor, Andrew Miri, and members of the Dombeck Lab for helpful comments and discussion on the manuscript. This work was supported by the NIH (NIMH R01-MH101297), a Hartwell Fellowship from the Hartwell Foundation (to JBI), and a NARSAD Young Investigator Grant from the Brain & Behavior Research Foundation (to JBI). rAAV2-Retro-CAG-TdTomato was a gift from the Janelia Research Campus and the GENIE project.

## Supplementary Figures

**Supplementary Figure 1:**
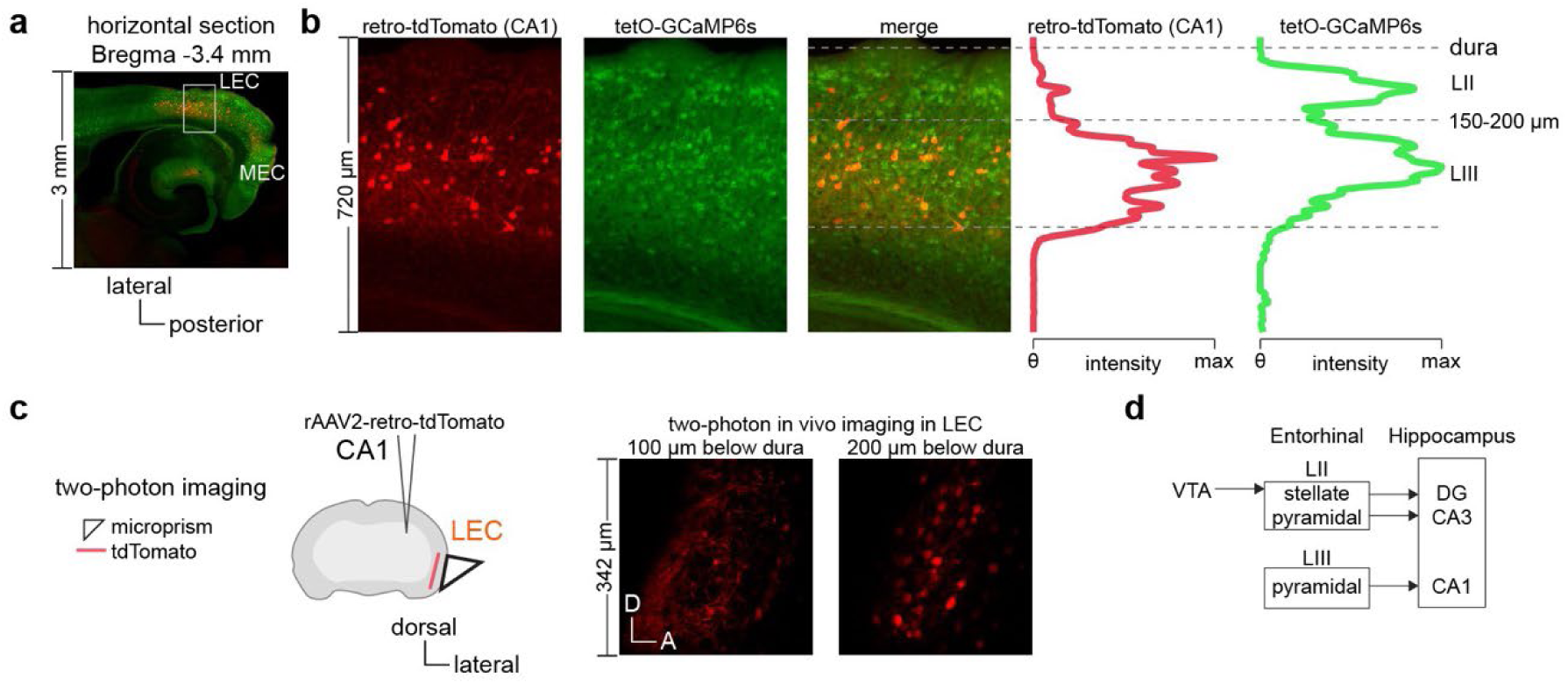
Histological confirmation of imaging window over LEC. a) Histology of lateral entorhinal cortex (LEC). Injection of retrograde tracer in CA1 labels CA1-projecting LIII pyramidal cells of entorhinal cortex in a mouse with GCaMP6s expression. Horizontal section is taken after PFA fixation. b) Zoomed in images of tdTomato and GCaMP6s labeling of the LEC in this horizontal slice. Histogram quantifies the amount of fluorescence from each of these channels as a function of depth from the dural surface. The cell-free zone separating layers II and III is approximately 150 to 200 µm beneath dura. c) To validate the prism placement, CA1-projecting LIII cells in LEC were again labeled with a retrograde tracer and two-photon imaging was performed on an implanted mouse. Labeled cell bodies were only seen at depths of at least 200 µm. d) Simplified diagram of connectivity between superficial layers of entorhinal cortex and regions of the hippocampus along with dopaminergic inputs from the ventral tegmental area (VTA).

**Supplementary Figure 2:**
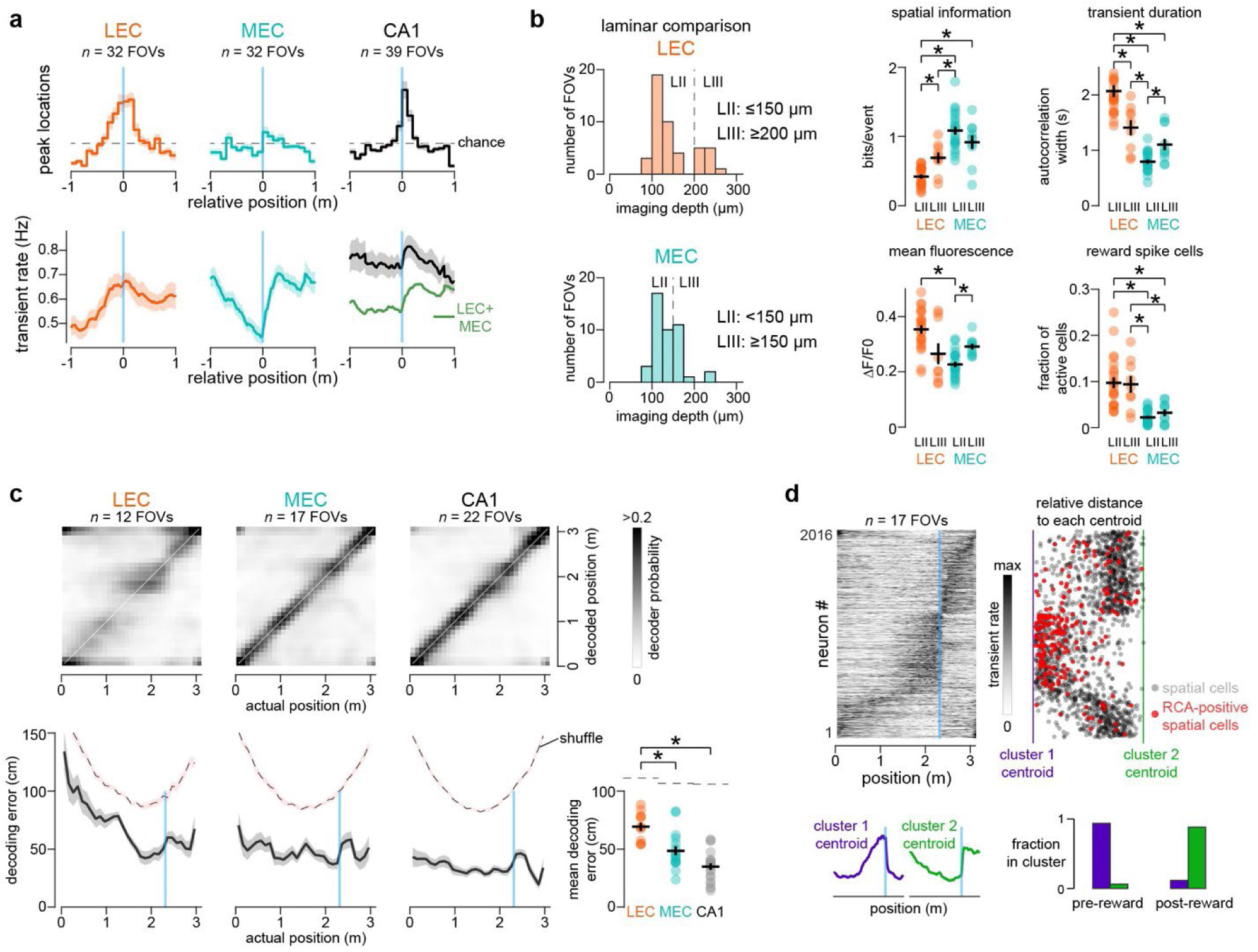
LEC firing peaks around reward and differs by layer. a) Histogram of spatial cell peaks and mean transient rates relative to the reward location. Datasets were combined across days where the reward location was either at 2.3 m (as in Figure 2a-d), 0.7 m, or 1.5 m. Histograms are binned every 10 cm; transient rate is sampled every 1 cm. Mean ± SEM are shown as dark line with light shading, respectively. Green LEC+MEC trace is the average of the LEC and MEC curves. b) Comparison of Ca^2+^ transients and cell firing properties from LII (23 fields) and LIII (9 fields) of the LEC and LII (22 fields) and LIII (10 fields) of the MEC. Data quantified for each imaging session; black cross represents mean ± SEM across sessions. Statistical tests performed compared between pairs (2-sample t-test, Bonferroni correction). * indicates p < 0.05 and n.s. indicates p > 0.05. Non-significant pairwise comparisons are not shown. Imaging depths for all LEC and all MEC fields (47 and 44 fields total, respectively, which include some sessions with tasks not included here) shown on left. c) Bayesian decoding of spatial position for sessions with the reward location at 2.3 m. A random subset of 40 non-reward spatial cells from each field of view was used to train a decoder (odd laps) and tested on the remaining laps (even laps). Because not all imaging fields contained at least 40 non-reward spatial cells, we only used data for the sessions with enough cells. Data was binned every 10 cm. In each bin, the decoded error was taken as the mean absolute difference between the most likely decoded position and the actual position. * indicates p < 0.05 using 2-sample t-test. Shuffle (shown as dashed lines) was achieved by randomly permuting the position data for the test set (even laps) and performing the same decoding analysis. d) Unsupervised k-means clustering of spatial firing patterns for LEC with reward location at 2.3 m (data shown in Figure 2c). Using the Calinski-Harabasz criterion, the optimal number of clusters was identified as 2. The cluster centroids are shown in purple and green, along with the fraction of pre-reward cells and post-reward cells that identify as each cluster. Dot raster shows relative distance to each centroid for all spatial cells (gray) or spatial cells that are also RCA cells (red), taken as (d_2_-d_1_)/(d_2_+d_1_), where d_1_ is the squared Euclidean distance to centroid 1 and d_2_ to centroid 2.

**Supplementary Figure 3:**
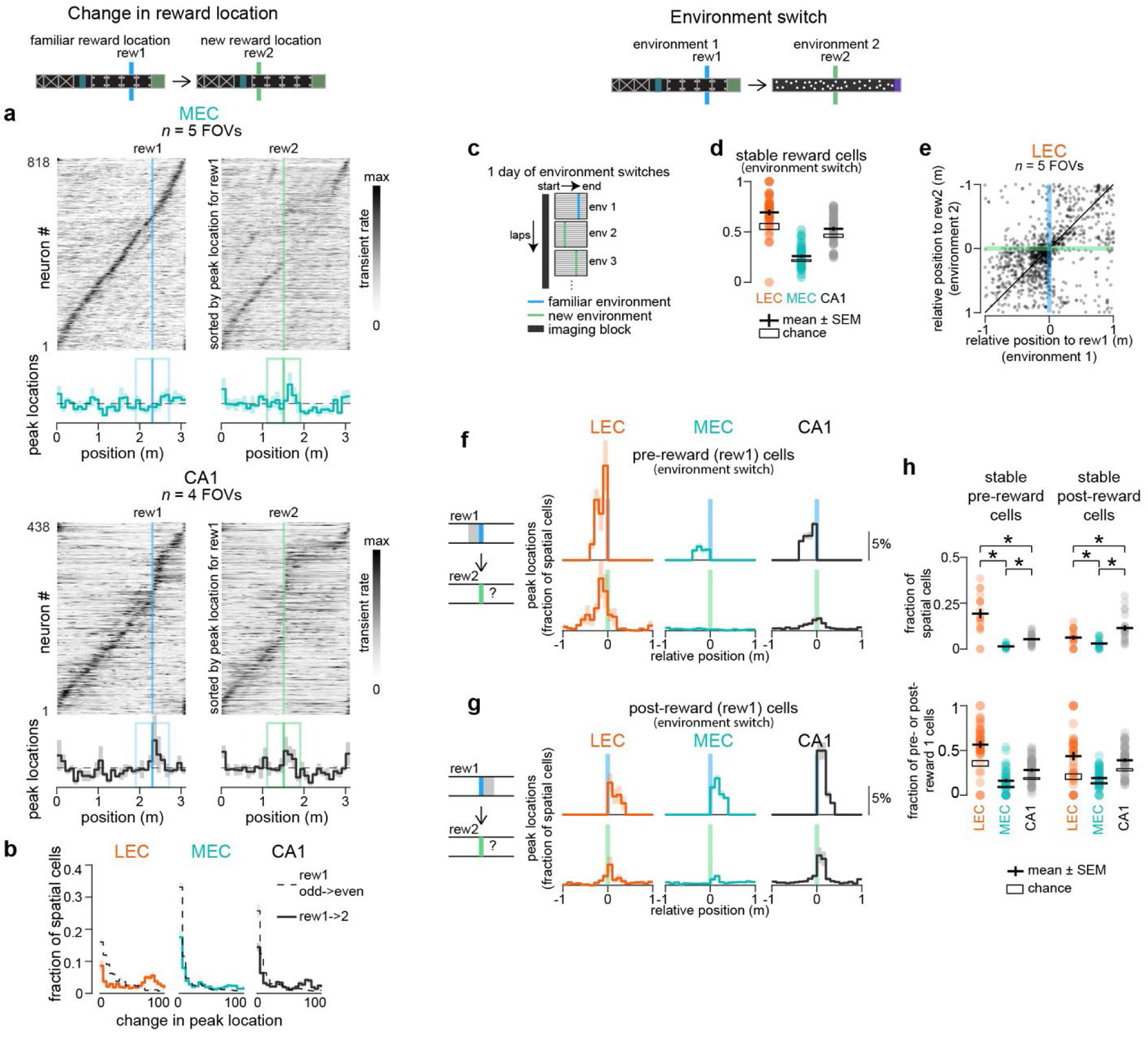
Additional details on dedication or pre- and post-reward populations for both reward moves and environment switches. a) Spatial firing patterns along the track and histogram of firing peaks for spatial cells sorted by their firing peaks with the familiar reward location for MEC and CA1. Same format as Figure 3c. b) Distribution of difference in peak location for MEC and CA1 spatial cells across the two conditions. Dashed line is the distribution for cross-validated data with the familiar reward location (even versus odd laps). c) Paradigm for environment switches. Four different environments were visited in the same session, with ~20 laps run in each environment before switching to the next one. d) The fraction of reward cells that remain reward cells after the environment is changed, quantified by the fraction of reward cells that maintain their peak location within 40 cm of the new reward location. Data quantified for each environment switch in an imaging session; black cross represents mean ± SEM across switches. e) Peak locations shown relative to reward locations with first environment on the x-axis and second environment on the y-axis. Each point represents one LEC neuron that was a spatial cell in both. f) Peak locations for pre-reward cells relative to reward locations in the two environments. Selected cells are chosen for being pre-reward cells with the first environment (rew1) and spatial cells in both; what is shown here is where their peak locations are for the second environment’s reward location (rew2). Histograms are calculated for each environment switch in an imaging session; mean ± SEM are shown as dark line with light shading, respectively. g) Same as panel f but for post-reward cells. h) Histograms quantify cells that are stable pre- or post-reward cells in both environments, either as a fraction of all spatial cells or as a fraction of the pre- or post-reward cells for the first environment. Each point represents one environment switch in an imaging session; black cross represents mean ± SEM; black rectangle represents mean ± SEM expected by chance. Statistical tests performed between each pair (2-sample t-test). * indicates p < 0.05 and n.s. indicates p > 0.05.

**Supplementary Figure 4:**
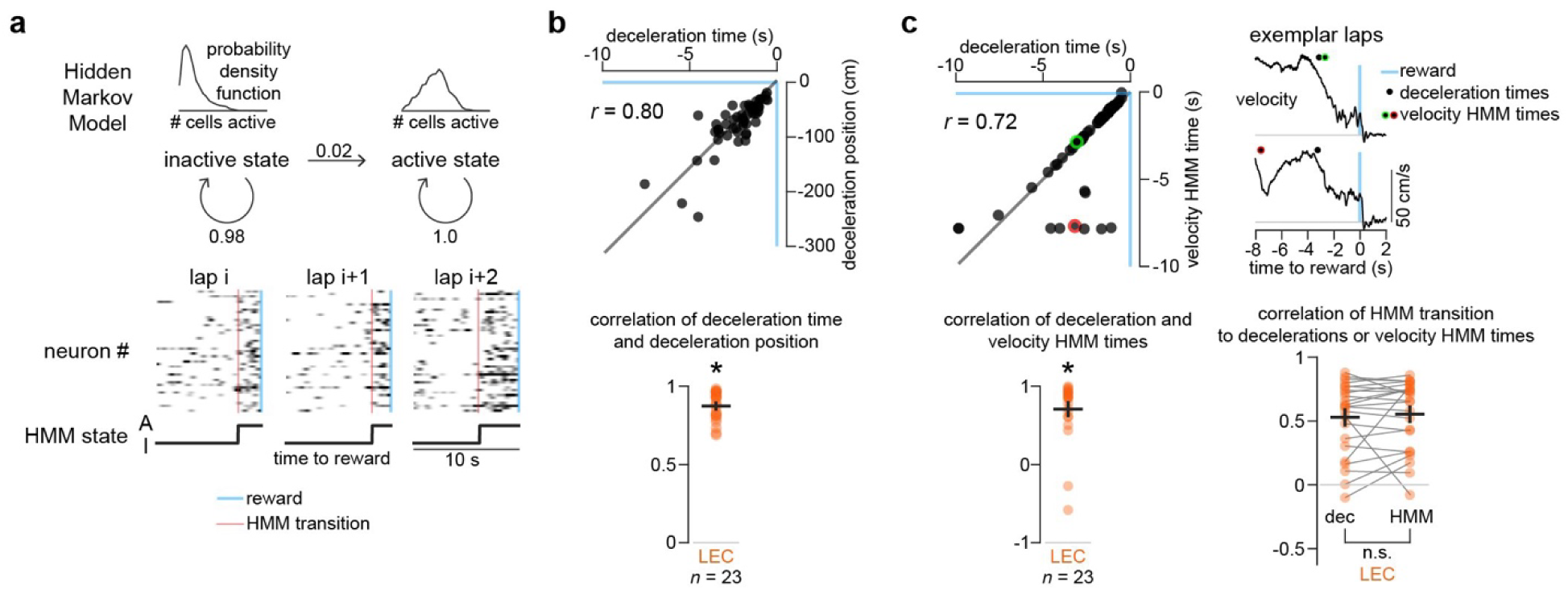
Additional details on hidden Markov model used to model LEC pre-reward population firing. a) A hidden Markov model (HMM) was used to detect transitions in the pre-reward population activity. The 10 seconds before reward delivery were considered. The system begins in an inactive state and can transition to an absorbing active state on each lap. The number of active cells is the observed data. The HMM is trained to learn the emission probabilities for each state (number of cells active) and the transition probability matrix. b) Deceleration times and position relative to reward are highly correlated, thus indicating that the running velocity is fairly consistent during the period of approach to reward. The analysis in the main text uses time because it was more reliable in practice. c) Transition from high running speed to lower running speed during reward approach was measured in two ways: deceleration time and velocity HMM time (see Methods). For exemplar session (top left: points represent transition times for individual laps; top right: exemplar laps where velocity HMM time agreed or disagreed with the deceleration time, indicated by green or red highlight, respectively) and across sessions (bottom left: each point represents correlation between the two measures for one imaging session; black cross represents mean ± SEM across sessions), the two measures were highly correlated. Both measures were also similarly highly correlated to the HMM-detected state change in the pre-reward cell firing (bottom right: Pearson correlation coefficients for each session, calculated as in Figure 4g; each point represents one imaging session; black cross represents mean ± SEM across sessions). Statistical tests performed against zero (1-sample t-test; * indicates p < 0.05) and between each pair of regions (Wilcoxon signed-rank test; n.s. indicates p > 0.05).

**Supplementary Figure 5:**
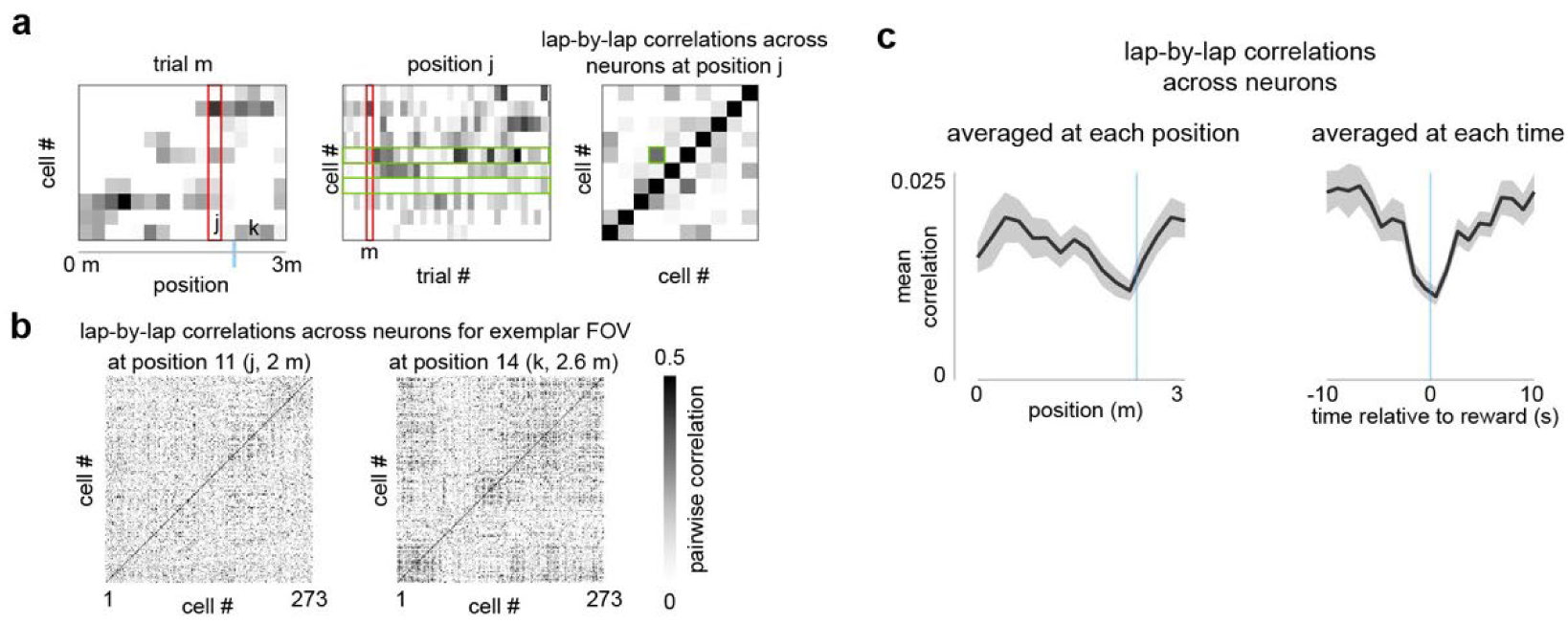
Pairwise correlations between LEC neurons across trials decrease near rewards. a) Correlation matrices were formed for the activity patterns of a neural population in a given imaging session at a given position. We calculated the correlation between the activity of each pair of cells at a given position on the track across all the laps for that session. b) Example correlation matrices at two track positions are shown. c) Correlations as a function of track position or time relative to reward. At a given position (or time), the correlation matrix was averaged across all cell-cell pairs (excluding the diagonal) and computed for each imaging session.

**Supplementary Figure 6:**
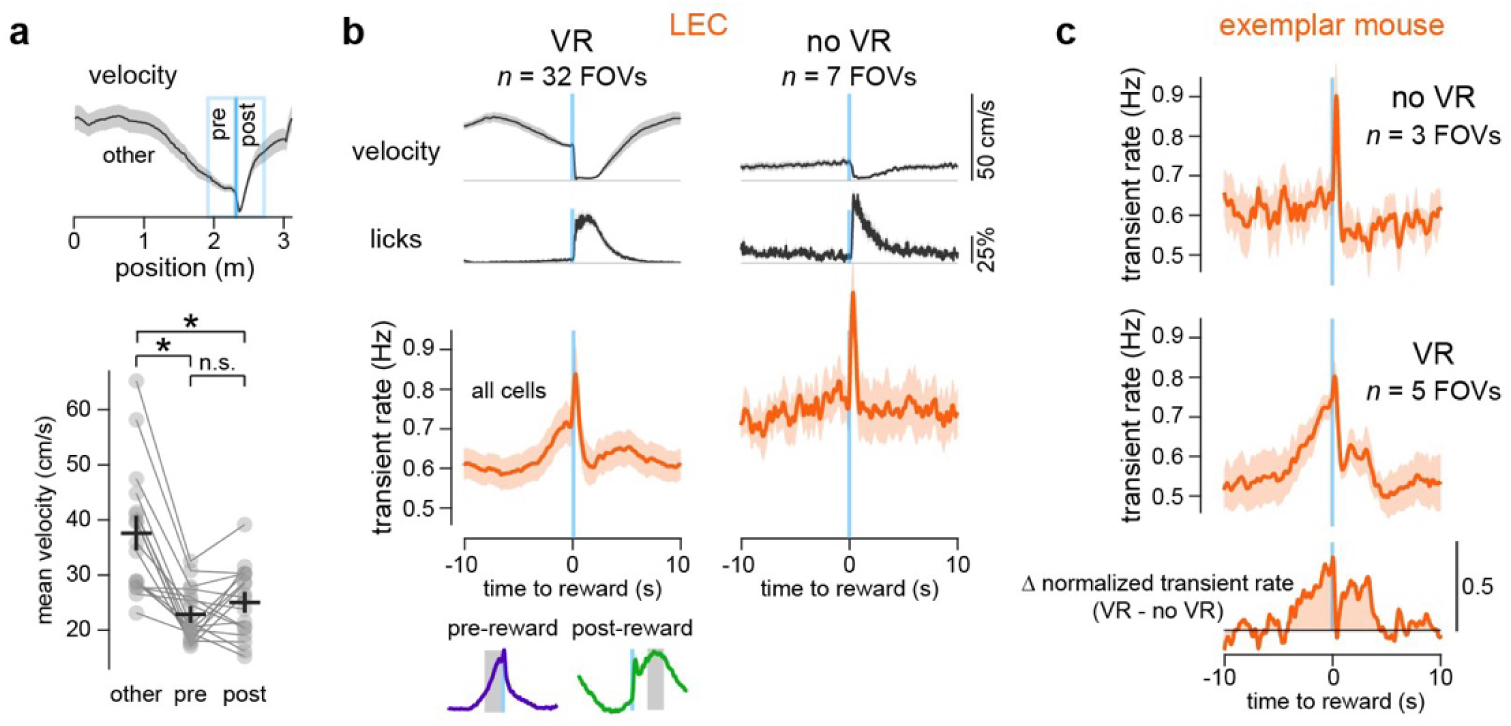
Pre- and post-reward firing in LEC is specific to a virtual navigation task. a) Mean running velocity at track positions surrounding reward are similar (pre: 40 cm before reward, mean of 22.8 cm/s; post: 40 cm after reward, mean of 25.0 cm/s) but lower than running speed along the rest of the track (other: all track positions excluding 40 cm around reward, mean of 37.6 cm/s). Each point represents one imaging session in LEC for reward at 2.3 m; black cross represents mean ± SEM across sessions. Statistical tests performed between each group (Wilcoxon signed-rank test; * indicates p < 0.05 and n.s. indicates p > 0.05). b) VR (left column): mean velocity, lick rate, and transient rate across all active cells in LEC, as a function of time relative to reward. LEC transient rate plot is reproduced from Figure 5a. Inset shows mean transient rate for pre-reward neurons and post-reward neurons relative to reward time. Gray boxes highlight periods when pre-reward and post-reward cells are highly active outside of the immediate reward consumption period. No VR (right column): same measures but for naïve head-fixed mice (7 FOVs in 3 mice) receiving random (unpredictable) rewards. We only included rewards with immediate consumption (first lick within 0.5 s of reward delivery) and with some treadmill movement in the 2.5 s preceding reward. c) In one exemplar mouse, after imaging LEC during delivery of random rewards (‘no VR’ condition), the same mouse was trained in virtual reality over a few days and imaging was performed during a reward navigation task in virtual reality (‘VR’ condition). Transient rate was averaged across all active neurons for each session. Mean ± SEM are shown as dark line with light shading, respectively. After normalizing transient rate (by dividing by the mean transient rate in the period from −10 s to −5 s), the difference in bulk firing between ‘VR’ and ‘no VR’ conditions was calculated. This difference curve shows two periods of large difference: pre-reward and post-reward, relieved by a brief moment of equality for the 0.5-1 s immediately after reward delivery.

**Supplementary Figure 7:**
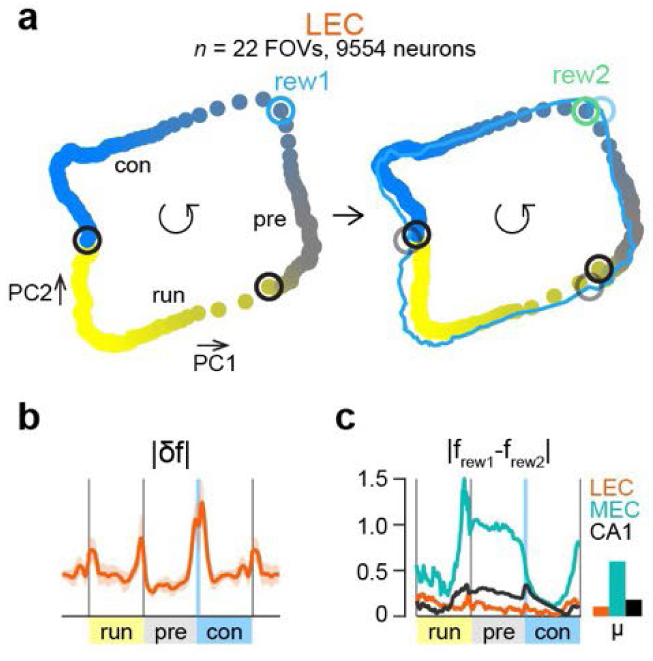
Trajectory of LEC population firing using state space analysis. a) Population LEC firing with respect to behavioral epochs is reduced to two dimensions using principal component analysis. The mean trajectory is plotted, with the color coded by the behavioral epoch (yellow: post-reward/running, gray: reward approach, blue: reward consumption). Open circles highlight the points in the trajectory that correspond to the transition points between epochs. The same components are used to plot the population firing for familiar reward (rew1) and novel reward (rew2). The trajectory for familiar reward is reproduced on the right plot using a thin blue line for comparison. b) Magnitude of the difference between successive points (1-norm of the derivative) using the first 10 principal components. c) Quantification of the difference between the trajectories for familiar and novel rewards using the first two principal component dimensions, taken as the 2-norm of the difference normalized by the 2-norm of the familiar reward trajectory.

---

**Supplementary Table 1.**
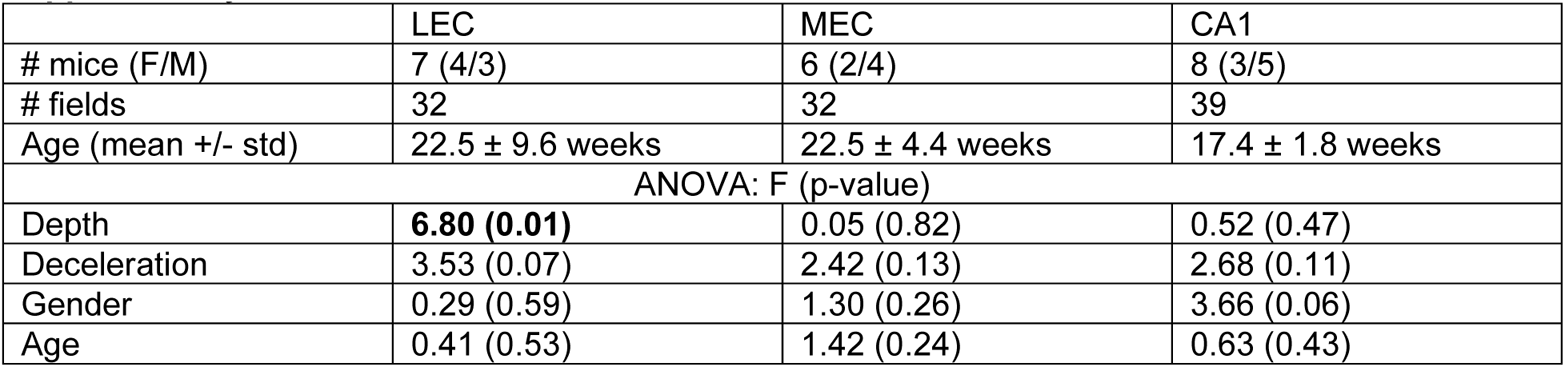
Multiway analysis of variance (ANOVA) was performed using the anovan function in MATLAB. The effects of the following factors were tested for their effect on reward clustering for each imaging session: 1) imaging depth, discretized into 50 µm bins; 2) degree of anticipatory deceleration, measured as mean velocity in 1 s before reward dropping to at least half of the maximum velocity; 3) gender, 4) age, in weeks.

|  | LEC | MEC | CA1 |
| --- | --- | --- | --- |
| # mice (F/M) | 7 (4/3) | 6 (2/4) | 8 (3/5) |
| # fields | 32 | 32 | 39 |
| Age (mean +/- std) | 22.5 ± 9.6 weeks | 22.5 ± 4.4 weeks | 17.4 ± 1.8 weeks |
| ANOVA: F (p-value) |  |  |  |
| Depth | **6.80 (0.01)** | 0.05 (0.82) | 0.52 (0.47) |
| Deceleration | 3.53 (0.07) | 2.42 (0.13) | 2.68 (0.11) |
| Gender | 0.29 (0.59) | 1.30 (0.26) | 3.66 (0.06) |
| Age | 0.41 (0.53) | 1.42 (0.24) | 0.63 (0.43) |

**Supplementary Table 2.** Number of LEC imaging fields and number of neurons for each of the three reward locations used. Final row shows data for sessions where the reward location was changed. For this data, ‘Spatial neurons’ shows number of spatial neurons for familiar and novel reward location, respectively.

| Reward location | # FOVs | Total neurons | Active neurons | Spatial neurons |
| --- | --- | --- | --- | --- |
| 0.7 m | 5 | 2106 | 2001 | 577 |
| 1.5 m | 10 | 4812 | 4387 | 1363 |
| 2.3 m | 17 | 8538 | 8101 | 2016 |
| Total (familiar reward) | 32 | 15456 | 14489 | 3956 |
| Total (reward change) | 22 | 10208 | 9554 | 2890, 2447 |

**Supplementary Table 3.** Number of sessions for optogenetic inhibition experiments in LEC.

|  | control | Jaws |
| --- | --- | --- |
| # mice (F/M) | 4 (1/3) | 5 (0/5) |
| # sessions | 25 | 26 |
| Age (mean +/- std) | 25.3 ± 2.8 weeks | 23.3 ± 1.7 weeks |

## References

1. Sosa, M. & Giocomo, L. M. Navigating for reward. Nat. Rev. Neurosci. 22, 472–487 (2021).

2. Nyberg, N., Duvelle, É., Barry, C. & Spiers, H. J. Spatial goal coding in the hippocampal formation. Neuron 110, 394–422 (2022).

3. Eichenbaum, H. Elements of Information Processing in Hippocampal Neuronal Activity: Space, Time, and Memory. in The Hippocampus from Cells to Systems: Structure, Connectivity, and Functional Contributions to Memory and Flexible Cognition (eds. Hannula, D. E. & Duff, M. C.) 69–94 (Springer International Publishing, 2017). doi:10.1007/978-3-319-50406-3_3.

4. Van Cauter, T. et al. Distinct Roles of Medial and Lateral Entorhinal Cortex in Spatial Cognition. Cereb. Cortex 23, 451–459 (2013).

5. Foster, D. J., Morris, R. G. & Dayan, P. A model of hippocampally dependent navigation, using the temporal difference learning rule. Hippocampus 10, 1–16 (2000).

6. Dayan, P. Improving Generalization for Temporal Difference Learning: The Successor Representation. Neural Comput. 5, 613–624 (1993).

7. Stachenfeld, K. L., Botvinick, M. M. & Gershman, S. J. The hippocampus as a predictive map. Nat. Neurosci. 20, 1643–1653 (2017).

8. Tulving, E. Elements of Episodic Memory. (Oxford University Press, 1983).

9. Eichenbaum, H., Dudchenko, P., Wood, E., Shapiro, M. & Tanila, H. The Hippocampus, Memory, and Place Cells: Is It Spatial Memory or a Memory Space? Neuron 23, 209–226 (1999).

10. Smith, D. M. & Mizumori, S. J. Y. Hippocampal place cells, context, and episodic memory. Hippocampus 16, 716–729 (2006).

11. Wood, E. R., Dudchenko, P. A., Robitsek, R. J. & Eichenbaum, H. Hippocampal Neurons Encode Information about Different Types of Memory Episodes Occurring in the Same Location. Neuron 27, 623–633 (2000).

12. Frank, L. M., Brown, E. N. & Wilson, M. Trajectory Encoding in the Hippocampus and Entorhinal Cortex. Neuron 27, 169–178 (2000).

13. Moita, M. A. P., Rosis, S., Zhou, Y., LeDoux, J. E. & Blair, H. T. Hippocampal Place Cells Acquire Location-Specific Responses to the Conditioned Stimulus during Auditory Fear Conditioning. Neuron 37, 485–497 (2003).

14. Markus, E. J. et al. Interactions between location and task affect the spatial and directional firing of hippocampal neurons. J. Neurosci. 15, 7079–7094 (1995).

15. O’Keefe, J. Place units in the hippocampus of the freely moving rat. Exp. Neurol. 51, 78–109 (1976).

16. Hollup, S. A., Molden, S., Donnett, J. G., Moser, M.-B. & Moser, E. I. Accumulation of Hippocampal Place Fields at the Goal Location in an Annular Watermaze Task. J. Neurosci. 21, 1635–1644 (2001).

17. Gauthier, J. L. & Tank, D. W. A Dedicated Population for Reward Coding in the Hippocampus. Neuron 99, 179–193.e7 (2018).

18. Eichenbaum, H., Kuperstein, M., Fagan, A. & Nagode, J. Cue-sampling and goal-approach correlates of hippocampal unit activity in rats performing an odor-discrimination task. J. Neurosci. 7, 716–732 (1987).

19. Aoki, Y., Igata, H., Ikegaya, Y. & Sasaki, T. The Integration of Goal-Directed Signals onto Spatial Maps of Hippocampal Place Cells. Cell Rep. 27, 1516–1527.e5 (2019).

20. Ranck, J. B. Studies on single neurons in dorsal hippocampal formation and septum in unrestrained rats: Part I. Behavioral correlates and firing repertoires. Exp. Neurol. 41, 462–531 (1973).

21. Nilssen, E. S., Doan, T. P., Nigro, M. J., Ohara, S. & Witter, M. P. Neurons and networks in the entorhinal cortex: A reappraisal of the lateral and medial entorhinal subdivisions mediating parallel cortical pathways. Hippocampus 29, 1238–1254 (2019).

22. Hafting, T., Fyhn, M., Molden, S., Moser, M.-B. & Moser, E. I. Microstructure of a spatial map in the entorhinal cortex. Nature 436, 801–806 (2005).

23. Solstad, T., Boccara, C. N., Kropff, E., Moser, M.-B. & Moser, E. I. Representation of Geometric Borders in the Entorhinal Cortex. Science 322, 1865–1868 (2008).

24. Giocomo, L. M. et al. Topography of Head Direction Cells in Medial Entorhinal Cortex. Curr. Biol. 24, 252–262 (2014).

25. Butler, W. N., Hardcastle, K. & Giocomo, L. M. Remembered reward locations restructure entorhinal spatial maps. Science 363, 1447–1452 (2019).

26. Boccara, C. N., Nardin, M., Stella, F., O’Neill, J. & Csicsvari, J. The entorhinal cognitive map is attracted to goals. Science 363, 1443–1447 (2019).

27. Grienberger, C. & Magee, J. C. Entorhinal cortex directs learning-related changes in CA1 representations. Nature 611, 554–562 (2022).

28. Krishnan, S., Heer, C., Cherian, C. & Sheffield, M. E. J. Reward expectation extinction restructures and degrades CA1 spatial maps through loss of a dopaminergic reward proximity signal. Nat. Commun. 13, 6662 (2022).

29. Kaufman, A. M., Geiller, T. & Losonczy, A. A Role for the Locus Coeruleus in Hippocampal CA1 Place Cell Reorganization during Spatial Reward Learning. Neuron 105, 1018–1026.e4 (2020).

30. Takeuchi, T. et al. Locus coeruleus and dopaminergic consolidation of everyday memory. Nature 537, 357–362 (2016).

31. Tsao, A., Moser, M.-B. & Moser, E. I. Traces of Experience in the Lateral Entorhinal Cortex. Curr. Biol. 23, 399–405 (2013).

32. Deshmukh, S. & Knierim, J. Representation of Non-Spatial and Spatial Information in the Lateral Entorhinal Cortex. Front. Behav. Neurosci. 5, (2011).

33. Tsao, A. et al. Integrating time from experience in the lateral entorhinal cortex. Nature 561, 57– 62 (2018).

34. Leitner, F. C. et al. Spatially segregated feedforward and feedback neurons support differential odor processing in the lateral entorhinal cortex. Nat. Neurosci. 19, 935–944 (2016).

35. Lee, J. Y. et al. Dopamine facilitates associative memory encoding in the entorhinal cortex. Nature 598, 321–326 (2021).

36. Tamamaki, N. & Nojyo, Y. Preservation of topography in the connections between the subiculum, field CA1, and the entorhinal cortex in rats. J. Comp. Neurol. 353, 379–390 (1995).

37. Mancini, M. et al. Head and Neck Veins of the Mouse. A Magnetic Resonance, Micro Computed Tomography and High Frequency Color Doppler Ultrasound Study. PLOS ONE 10, e0129912 (2015).

38. Wekselblatt, J. B., Flister, E. D., Piscopo, D. M. & Niell, C. M. Large-scale imaging of cortical dynamics during sensory perception and behavior. APSselect 3, 2852–2866 (2016).

39. Chen, T.-W. et al. Ultrasensitive fluorescent proteins for imaging neuronal activity. Nature 499, 295–300 (2013).

40. Pachitariu, M. et al. Suite2p: beyond 10,000 neurons with standard two-photon microscopy. 061507 Preprint at 10.1101/061507 (2017).

41. Heys, J. G., Rangarajan, K. V. & Dombeck, D. A. The Functional Micro-organization of Grid Cells Revealed by Cellular-Resolution Imaging. Neuron 84, 1079–1090 (2014).

42. Dombeck, D. A., Harvey, C. D., Tian, L., Looger, L. L. & Tank, D. W. Functional imaging of hippocampal place cells at cellular resolution during virtual navigation. Nat. Neurosci. 13, 1433–1440 (2010).

43. Climer, J. R. & Dombeck, D. A. Information theoretic approaches to deciphering the neural code with functional fluorescence imaging. eNeuro (2021) doi:10.1523/ENEURO.0266-21.2021.

44. Fallon, J. H., Koziell, D. A. & Moore, R. Y. Catecholamine innervation of the basal forebrain II. Amygdala, suprarhinal cortex and entorhinal cortex. J. Comp. Neurol. 180, 509–531 (1978).

45. Farrell, K., Lak, A. & Saleem, A. B. Midbrain dopamine neurons signal phasic and ramping reward prediction error during goal-directed navigation. Cell Rep. 41, 111470 (2022).

46. Basu, R. et al. The orbitofrontal cortex maps future navigational goals. Nature 599, 449–452 (2021).

47. Howe, M. W., Tierney, P. L., Sandberg, S. G., Phillips, P. E. M. & Graybiel, A. M. Prolonged dopamine signalling in striatum signals proximity and value of distant rewards. Nature 500, 575–579 (2013).

48. Lloyd, K. & Dayan, P. Tamping Ramping: Algorithmic, Implementational, and Computational Explanations of Phasic Dopamine Signals in the Accumbens. PLOS Comput. Biol. 11, e1004622 (2015).

49. Latimer, K. W., Yates, J. L., Meister, M. L. R., Huk, A. C. & Pillow, J. W. Single-trial spike trains in parietal cortex reveal discrete steps during decision-making. Science 349, 184–187 (2015).

50. Sheffield, M. E. J., Adoff, M. D. & Dombeck, D. A. Increased Prevalence of Calcium Transients across the Dendritic Arbor during Place Field Formation. Neuron 96, 490–504.e5 (2017).

51. Chuong, A. S. et al. Noninvasive optical inhibition with a red-shifted microbial rhodopsin. Nat. Neurosci. 17, 1123–1129 (2014).

52. Tolman, E. C. Cognitive maps in rats and men. Psychol. Rev. 55, 189–208 (1948).

53. Wilson, D. I. G. et al. Lateral entorhinal cortex is critical for novel object-context recognition. Hippocampus 23, 352–366 (2013).

54. Kuruvilla, M. V., Wilson, D. I. G. & Ainge, J. A. Lateral entorhinal cortex lesions impair both egocentric and allocentric object-place associations. Brain Neurosci. Adv. 4, 2398212820939463 (2020).

55. Wang, C. et al. Egocentric coding of external items in the lateral entorhinal cortex. Science 362, 945–949 (2018).

56. Knierim, J. J., Neunuebel, J. P. & Deshmukh, S. S. Functional correlates of the lateral and medial entorhinal cortex: objects, path integration and local–global reference frames. Philos. Trans. R. Soc. B Biol. Sci. 369, 20130369 (2014).

57. Morris, R. G. M., Garrud, P., Rawlins, J. N. P. & O’Keefe, J. Place navigation impaired in rats with hippocampal lesions. Nature 297, 681–683 (1982).

58. Heys, J. G., Wu, Z., Allegra Mascaro, A. L. & Dombeck, D. A. Inactivation of the Medial Entorhinal Cortex Selectively Disrupts Learning of Interval Timing. Cell Rep. 32, 108163 (2020).

59. Lisman, J. E. & Grace, A. A. The Hippocampal-VTA Loop: Controlling the Entry of Information into Long-Term Memory. Neuron 46, 703–713 (2005).

60. Glovaci, I. & Chapman, C. A. Dopamine induces release of calcium from internal stores in layer II lateral entorhinal cortex fan cells. Cell Calcium 80, 103–111 (2019).

61. Stephens, D. W. & Krebs, J. R. Foraging Theory. (Princeton University Press, 1986).

62. Pirolli, P. & Card, S. Information foraging. Psychol. Rev. 106, 643–675 (1999).

63. Averbeck, B. B., Latham, P. E. & Pouget, A. Neural correlations, population coding and computation. Nat. Rev. Neurosci. 7, 358–366 (2006).

64. Panzeri, S., Moroni, M., Safaai, H. & Harvey, C. D. The structures and functions of correlations in neural population codes. Nat. Rev. Neurosci. 23, 551–567 (2022).

65. Eichenbaum, H. & Fortin, N. J. The neurobiology of memory based predictions. Philos. Trans. R. Soc. B Biol. Sci. 364, 1183–1191 (2009).

66. Schultz, W. Dopamine signals for reward value and risk: basic and recent data. Behav. Brain Funct. 6, 24 (2010).

67. Masurkar, A. V. et al. Medial and Lateral Entorhinal Cortex Differentially Excite Deep versus Superficial CA1 Pyramidal Neurons. Cell Rep. 18, 148–160 (2017).

68. Bittner, K. C., Milstein, A. D., Grienberger, C., Romani, S. & Magee, J. C. Behavioral time scale synaptic plasticity underlies CA1 place fields. Science 357, 1033–1036 (2017).

69. Sheffield, M. E. J. & Dombeck, D. A. Calcium transient prevalence across the dendritic arbour predicts place field properties. Nature 517, 200–204 (2015).

70. Yonelinas, A. P., Ranganath, C., Ekstrom, A. D. & Wiltgen, B. J. A contextual binding theory of episodic memory: systems consolidation reconsidered. Nat. Rev. Neurosci. 20, 364–375 (2019).

71. Gruber, M. J., Ritchey, M., Wang, S.-F., Doss, M. K. & Ranganath, C. Post-learning Hippocampal Dynamics Promote Preferential Retention of Rewarding Events. Neuron 89, 1110–1120 (2016).

72. Knierim, J. J., Lee, I. & Hargreaves, E. L. Hippocampal place cells: Parallel input streams, subregional processing, and implications for episodic memory. Hippocampus 16, 755–764 (2006).

73. Lisman, J. E. Role of the dual entorhinal inputs to hippocampus: a hypothesis based on cue/action (non-self/self) couplets. Prog. Brain Res. 163, 615–625 (2007).

74. Soma, S. et al. Rat hippocampal CA1 region represents learning-related action and reward events with shorter latency than the lateral entorhinal cortex. *Commun*. Biol. 6, 1–14 (2023).

75. Kim, H. R. et al. A Unified Framework for Dopamine Signals across Timescales. Cell 183, 1600–1616.e25 (2020).

76. Bowler, J. C. & Losonczy, A. Direct Cortical Inputs to Hippocampal Area CA1 Transmit Complementary Signals for Goal-directed Navigation. 2022.11.10.516009 Preprint at 10.1101/2022.11.10.516009 (2022).

77. Witter, M. P. & Moser, E. I. Spatial representation and the architecture of the entorhinal cortex. Trends Neurosci. 29, 671–678 (2006).

## Methods References

1. Leitner, F. C. et al. Spatially segregated feedforward and feedback neurons support differential odor processing in the lateral entorhinal cortex. Nature Neuroscience 19, 935–944 (2016).

2. Livneh, Y. et al. Homeostatic circuits selectively gate food cue responses in insular cortex. Nature 546, 611–616 (2017).

3. Heys, J. G., Rangarajan, K. V. & Dombeck, D. A. The Functional Micro-organization of Grid Cells Revealed by Cellular-Resolution Imaging. Neuron 84, 1079–1090 (2014).

4. Chia, T. H. & Levene, M. J. Microprisms for In Vivo Multilayer Cortical Imaging. Journal of Neurophysiology 102, 1310–1314 (2009).

5. van Groen, T. Entorhinal cortex of the mouse: Cytoarchitectonical organization. Hippocampus 11, 397–407 (2001).

6. Chuong, A. S. et al. Noninvasive optical inhibition with a red-shifted microbial rhodopsin. Nat Neurosci 17, 1123–1129 (2014).

7. Pisanello, F. et al. Dynamic illumination of spatially restricted or large brain volumes via a single tapered optical fiber. Nat Neurosci 20, 1180–1188 (2017).

8. Aronov, D. & Tank, D. W. Engagement of Neural Circuits Underlying 2D Spatial Navigation in a Rodent Virtual Reality System. Neuron 84, 442–456 (2014).

9. Guizar-Sicairos, M., Thurman, S. T. & Fienup, J. R. Efficient subpixel image registration algorithms. *Opt. Lett.*, OL 33, 156–158 (2008).

10. Pachitariu, M. et al. Suite2p: beyond 10,000 neurons with standard two-photon microscopy. 061507 Preprint at 10.1101/061507 (2017).

11. Friedrich, J., Zhou, P. & Paninski, L. Fast online deconvolution of calcium imaging data. PLOS Computational Biology 13, e1005423 (2017).

12. Huang, L. et al. Relationship between simultaneously recorded spiking activity and fluorescence signal in GCaMP6 transgenic mice. eLife 10, e51675 (2021).

13. Chen, T.-W. et al. Ultrasensitive fluorescent proteins for imaging neuronal activity. Nature 499, 295–300 (2013).

14. Smetters, D., Majewska, A. & Yuste, R. Detecting Action Potentials in Neuronal Populations with Calcium Imaging. Methods 18, 215–221 (1999).

15. Éltes, T., Szoboszlay, M., Kerti-Szigeti, K. & Nusser, Z. Improved spike inference accuracy by estimating the peak amplitude of unitary [Ca2+] transients in weakly GCaMP6f-expressing hippocampal pyramidal cells. The Journal of Physiology 597, 2925–2947 (2019).

16. Climer, J. R. & Dombeck, D. A. Information theoretic approaches to deciphering the neural code with functional fluorescence imaging. eNeuro (2021) doi:10.1523/ENEURO.0266-21.2021.

17. Helmchen, F., Imoto, K. & Sakmann, B. Ca2+ buffering and action potential-evoked Ca2+ signaling in dendrites of pyramidal neurons. Biophysical Journal 70, 1069–1081 (1996).

18. Etter, G., Manseau, F. & Williams, S. A Probabilistic Framework for Decoding Behavior From in vivo Calcium Imaging Data. Front Neural Circuits 14, 19 (2020).

